# High-fat diet enhances starvation-induced hyperactivity via sensitizing hunger-sensing neurons in *Drosophila*

**DOI:** 10.1101/612234

**Authors:** Rui Huang, Tingting Song, Haifeng Su, Liming Wang

**Affiliations:** Medical school, Chongqing University, Chongqing, 400030, China; Key Laboratory for Biorheological Science and Technology of Ministry of Education, State and Local Joint Engineering Laboratory for Vascular Implants, Bioengineering College, Chongqing University, Chongqing, 400030, China; MOE Key Laboratory of Biosystems Homeostasis & Protection and Innovation Center for Cell Signaling Network, Life Sciences Institute, Zhejiang University, Hangzhou, 310058, Zhejiang, China.; Institute of Neuroscience, State Key Laboratory of Neuroscience, CAS Center for Excellence in Brain Science and Intelligence Technology, Shanghai Institutes for Biological Sciences, Chinese Academy of Sciences, Shanghai, 200031, China.

**Author notes:** These authors contribute equally to this work (Rui Huang, Tingting Song, Haifeng Su).

## Abstract

Proper regulation of food intake can be disrupted by sustained metabolic challenges such as high-fat diet (HFD), which may result in various metabolic disorders. Previously, we showed that starvation induced sustained hyperactivity, an exploratory component of food-seeking behavior, via a specific group of octopamingeric (OA) neurons (Yu et al., 2016). In the present study, we found that HFD greatly enhanced starvation-induced hyperactivity. HFD increased the excitability of these OA neurons to a hunger hormone named adipokinetic hormone (AKH), via increasing the accumulation of AKH receptor (AKHR) in these neurons. Upon HFD, excess dietary lipids were transported by a lipoprotein LTP to enter these OA neurons via the cognate receptor LpR1, which activated AMPK-TOR signaling and suppressed autophagy-dependent degradation of AKHR. Taken together, we uncovered a mechanism that linked HFD and starvation-induced hyperactivity, providing insight in the reshaping of neural circuitry under metabolic challenges and the progression of metabolic diseases.

## INTRODUCTION

Obesity and obesity-associated metabolic disorders such as type 2 diabetes and cardiovascular diseases have become a global epidemic. Chronic over-nutrition, especially excessive intake of dietary lipids, is one of the leading causes of these metabolic disturbances (Bray and Popkin, 1998; Hill et al., 2000). Accumulating evidence has shown that HFD imposes adverse effects on the physiology and metabolism of liver, skeletal muscle, the adipose tissue, and the nervous system (Dietrich et al., 2013; Gomez-Perez et al., 2012; Liu et al., 2015; Shimobayashi et al., 2018). It is therefore of importance to understand the mechanisms underlying HFD-induced changes in different organs and cell types, which will offer critical insight into the diagnosis and treatment of obesity and other metabolic diseases.

The central nervous system plays a critical role in regulating food intake, energy homeostasis, and organismal metabolism (Gao and Horvath, 2007). In rodent models, neurons located in the arcuate nucleus of the hypothalamus, particularly neurons expressing Neuropeptide Y (NPY) and Agouti-Related Neuropeptide (AgRP) or those expressing Pro-opiomelanocortin (POMC), are important behavioral and metabolic regulators (Belgardt et al., 2009; Ollmann et al., 1997; Stanley and Leibowitz, 1984). These neurons detect various neural and hormonal cues such as circulating glucose and fatty acids, leptin, and ghrelin, and modulate food consumption and energy expenditure accordingly (Belgardt et al., 2009). Upon the reduction of the internal energy state, NPY/AgRP neurons are activated and exert a robust orexigenic effect (Belgardt et al., 2009). Genetic ablation of NPY/AgRP neurons in neonatal mice completely abolishes food consumption whereas acute activation of these neurons significantly enhances food consumption (Aponte et al., 2011; Krashes et al., 2011). NPY/AgRP neurons also antagonize the function of POMC neurons that plays a suppressive role on food consumption (Roseberry et al., 2004). Taken together, these two groups of neurons, among other neuronal populations, work in synergy to ensure a refined balance between energy intake and expenditure, and hence organismal metabolism.

In spite of their critical roles, the function of the nervous system to accurately regulate appetite and metabolism may be disrupted by sustained metabolic stress, resulting in eating disorders and various metabolic diseases such as obesity and type 2 diabetes. Several lines of evidence have begun to reveal the underlying neural mechanisms. For example, HFD increases the intrinsic excitability of orexigenic NPY/AgRP neurons (Vernia et al., 2016), induces leptin resistance (Mazor et al., 2018; Olofsson et al., 2013), and enhances their inhibitory innervations with anorexigenic POMC neurons (Newton et al., 2013), altogether resulting in hypersensitivity to starvation and increased food consumption. Interestingly, besides HFD, other metabolic challenges, including maternal HFD, alcohol consumption, as well as aging, also disrupt normal food intake via affecting the excitability and/or innervation of NPY/AgRP neurons (Cains et al., 2017; Furedi et al., 2018; Rivera et al., 2015). All these interventions may contribute to the onset and progression of metabolic disorders.

Proceeding the actual food consumption, food-seeking behavior is a critical yet largely overlooked behavioral component for the localization and occupation of desirable food sources. Food-seeking behavior has been characterized in rodent models, primarily by the elevation of locomotor activity and increased food approach of starved animals (Davidson, 2009; Mistlberger, 2011). It has been reported that NPY/AgRP neurons also play a role in food-seeking behavior (Aponte et al., 2011). However, to ensure adequate food intake, food seeking and food consumption are temporarily separated and even reciprocally inhibited (Betley et al., 2015; Chen et al., 2015). It remains largely unclear how the neural circuitry of food seeking and food consumption segregated and independently regulated in rodent models. Furthermore, it remains unknown whether HFD also affects food seeking, and if so whether its effects on both food seeking and food consumption share common mechanisms or not. To fully understand the intervention of energy homeostasis by sustained metabolic stress, we need to dissect the neural circuitry underlying food seeking and examine whether and how it is affected by HFD.

Fruit flies *Drosophila melanogaster* share fundamental analogy to vertebrate counterparts on the regulation of energy homeostasis and organismal metabolism despite that they diverged several hundred million years ago (Pandey and Nichols, 2011; Rajan and Perrimon, 2013; Reiter et al., 2001). Therefore, it offers a good model to characterize food-seeking behavior in depth and provide insight onto the regulation of energy intake and the pathogenesis of metabolic disorders in more complex organisms such as rodents and human.

Our previous work has shown that fruit flies exhibit robust starvation-induced hyperactivity that was directed towards the localization and acquisition of food sources, therefore resembling an important aspect of food-seeking behavior upon starvation (Yang et al., 2015). We have also identified a small subset of OA neurons in the fly brain that specifically regulate starvation-induced hyperactivity (Yu et al., 2016). Analogous to mammalian systems, a number of neural and hormonal cues are involved in the systemic control of nutrient metabolism and food intake. Among them, Neuropeptide F (NPF), short NPF (sNPF), Leucokinin, and Allatostatin A (AstA), have been shown to regulate food consumption, all of which have known mammalian homologs that regulate food intake (Pool and Scott, 2014). In particular, starvation-induced hyperactivity is regulated by two classes of neuroendocrine cells (Yu et al., 2016). One is functionally analogous to pancreatic α cells and produce AKH upon starvation, whereas the other produce *Drosophila* insulin-like peptides (DILPs), resembling the function of pancreatic β cells. These two classes of *Drosophila* hormones exert antagonistic functions on starvation-induced hyperactivity via the same group of OA neurons in the fly brain (Yu et al., 2016).

Based on these findings, we therefore sought to examine whether HFD disrupted the regulation of starvation-induced hyperactivity in fruit flies and aimed to investigate the underlying mechanism. In this present study, we found that HFD-fed flies rapidly became significantly more sensitive to starvation and exhibited starvation-induced hyperactivity much earlier than flies fed with normal food (ND). Meanwhile, HFD did not alter flies’ food consumption, suggesting that starvation-induced hyperactivity and food consumption are independently affected by HFD. Several days of HFD treatment did not alter the production of important hormonal cues like AKH and DILPs, but rather increased the sensitivity of the OA neurons that regulated starvation-induced hyperactivity to the hunger hormone AKH. In these OA neurons, constitutive autophagy maintained the level of AKHR protein, which determined their sensitivity to AKH and hence starvation. HFD feeding suppressed neuronal autophagy via AMPK-TOR signaling and in turn increased the level of AKHR in these OA neurons. Consistently, eliminating autophagy in these neurons mimicked the effect of HFD on starvation-induced hyperactivity whereas promoting autophagy inhibited the induction of hyperactivity by starvation. Furthermore, we also showed that a specific lipoprotein LTP and its cognate receptor LpR1 likely mediated the effect of HFD on the neuronal autophagy of OA neurons and hence its effect on starvation-induced hyperactivity. Taken together, we uncovered a novel mechanism that linked HFD, AMPK-TOR signaling, neuronal autophagy, and starvation-induced hyperactivity, shedding crucial light on the reshaping of neural circuitry under metabolic stress and the progression of metabolic diseases.

## RESULTS

### HFD specifically enhanced starvation-induced hyperactivity

We have previously reported that starvation induced hyperactivity in fruit flies *Drosophila melanogaster*, which facilitated the localization and acquisition of food and hence partly resembled food-seeking behavior (Yang et al., 2015; Yu et al., 2016). In this present study, we first asked whether HFD affected starvation-induced hyperactivity in fruit flies.

Flies were fed with normal fly food (ND) or HFD (by adding 20% coconut oil in the ND) for five consecutive days before tested in a locomotion assay that measured the frequency to cross the midline of tubes in the *Drosophila* Activity Monitor System (DAMS, Trikinetics) (Figure 1A). The midline-crossing frequency assayed by DAMS offered a reliable measure of flies’ locomotor activity and hence starvation-induced hyperactivity (Yang et al., 2015; Yu et al., 2016). ND-fed flies exhibited a robust increase in locomotion upon starvation (Figure 1A-B, blue, Day 1 and Day 2), which was consistent with our previous findings. Meanwhile, flies fed with HFD exhibited further enhanced hyperactivity upon starvation, while their baseline activity was comparable to ND-fed flies (Figure 1A-B, orange, Day 1 and Day 2). These results suggest that HFD-fed flies are behaviorally more sensitive to starvation.

**Figure 1.**
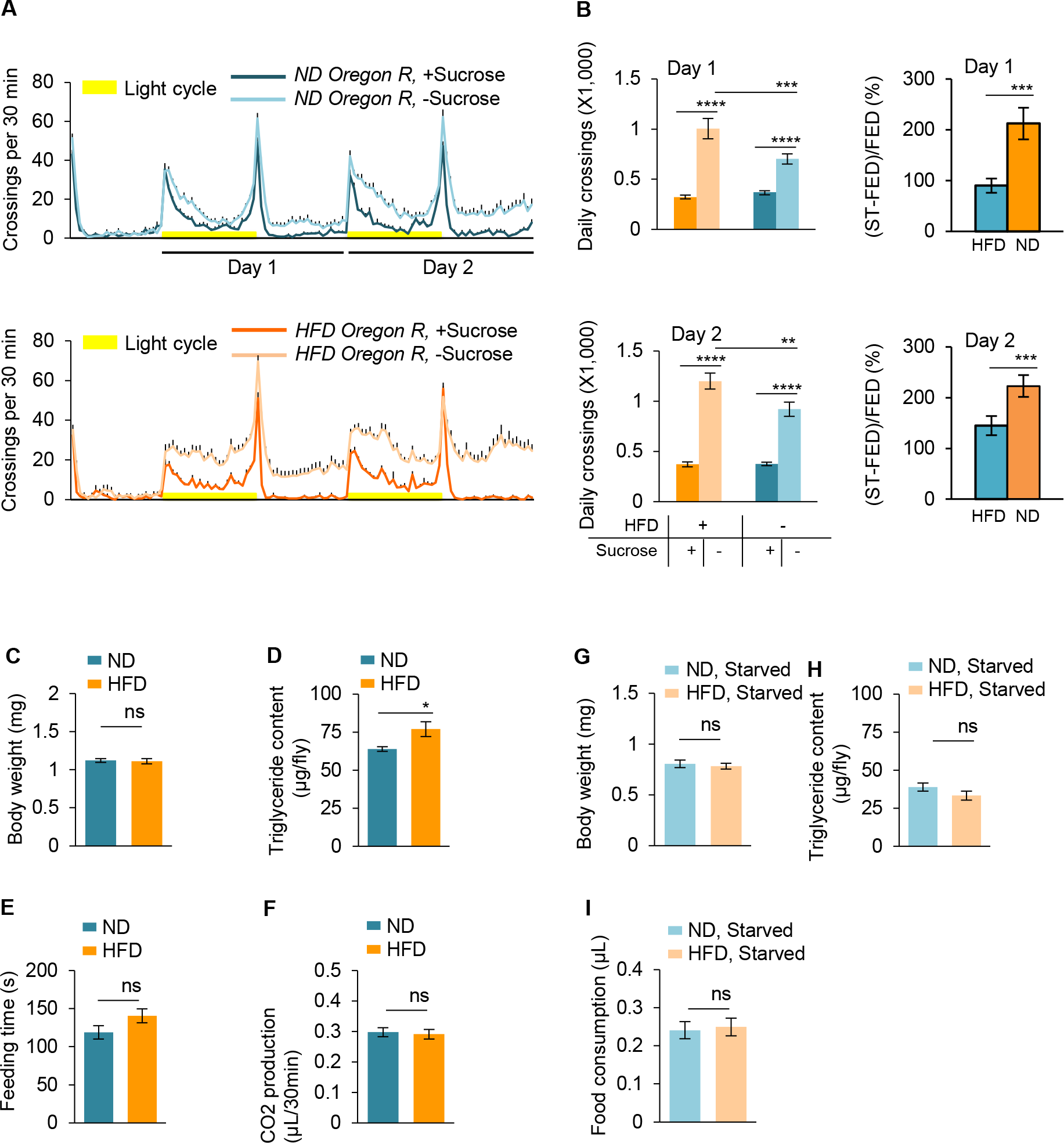
HFD promotes starvation-induced hyperactivity in adult *Drosophila*. (A) Wild-type *Oregon-R* virgin female flies fed with normal fly food (ND, blue) or high-fat diet (HFD, orange) were assayed in the presence (dark color) and absence (light color) of 5% sucrose using DAMS-based locomotion assay (n = 46-63). Midline crossing activity of indicated flies were shown. Yellow bars represent 12 hr light-on period in this and following figures. (B) Average daily midline crossing activity of flies assayed in A (left), and starvation-induced hyperactivity after 5-day HFD feeding vs. ND feeding experience (right). Data were broken down to Day 1 (upper) and Day 2 (lower). In the following figures only Day 1 data was presented. (C-D) Average body weight (C) and triglyceride storage (D) of indicated flies fed *ad libitum* (n = 6-18). (E) Total duration of feeding time during 24 hr recording in the FLIC assay (n = 38-47). (F) Total CO_2_ production for 1 hr of flies fed with ND or HFD (n = 12-14). (G-I) Average body weight (G, n = 18), triglyceride storage (H, n = 6), and 100 mM sucrose consumption (I, n = 30-35) of indicated flies starved for 36 hr (ST: starvation). ns, p > 0.05; *p < 0.05; ***p < 0.001; ****p < 0.0001. Student’s t-test and one-way ANOVA followed by post hoc test weith Bonferroni correction were used for pair-wise and multiple comparisons, respectively.

We then sought to understand how HFD feeding enhanced starvation-induced hyperactivity. One possible explanation was that HFD might decrease flies’ energy storage or increase flies’ energy expenditure, reducing their resistence to starvation. It seemed unlikely, since flies’ body weight was not affected by HFD and their lipid storage was actually slightly elevated by HFD (Figure 1C-D). Meanwhile, HFD-fed flies exhibited comparable feeding activity in the FLIC assay (Ro et al., 2014) as well as CO_2_ production compared to ND-fed flies, suggesting that their energy intake and expenditure remained unaffected (Figure 1E-F).

Upon starvation, ND-fed and HFD-fed flies exhibited comparable survival rate (Figure 1-figure supplement 1), suggesting that prior experience of HFD feeding did not alter starvation resistance. Consistently, upon starvation, the body weight and lipid storage of both ND-fed and HFD-fed flies decreased (Figure 1G-H) whereas the food consumption of these two group of flies increased in the MAFE assay in comparable manners (Figure 1I, note that fed flies did not exhibit food ingestion in the MAFE assays so we tested them in the FLIC assay, see Figure 1E) (Qi et al., 2015). Taken together, these results suggest that five days of HFD feeding specifically enhances the onset of starvation-induced hyperactivity, without altering flies’ resistence to starvation or food-consumption behavior.

### HFD increased the sensitivity of hunger-sensing, AKHR^+^ neurons

We have shown in a previous study that starvation-induced hyperactivity of adult flies was driven by a specific group of OA neurons in the fly brain (Yu et al., 2016). As a hunger sensor, the activity of these OA^+^ neurons is regulated by two groups of functionally antagonizing hormones, the hunger-induced hormone AKH and the satiety hormones DILPs. Therefore, we reasoned that HFD might enhance starvation-induced hyperactivity via modulating the signaling of these two groups of hormones.

We first tested whether the expression of AKH and DILP2 (the major DILP expressed in the fly brain) was modulated by HFD feeding. As shown in Figure 2A-B, ND-fed and HFD-fed flies showed comparable AKH and DILP2 mRNA production as revealed by both RNAseq (Figure 2A) and quantitative RT-PCR (Figure 2B) using head tissues. Moreover, circulating AKH and DILP2 in flies’ hemolymph remained unaffected by HFD feeding, as shown by the dot blot using antibodies against AKH and DILP2 proteins (Figure 2C). Therefore, the production of these hormones that regulated starvation-induced hyperactivity was not regulated by HFD feeding.

**Figure 2.**
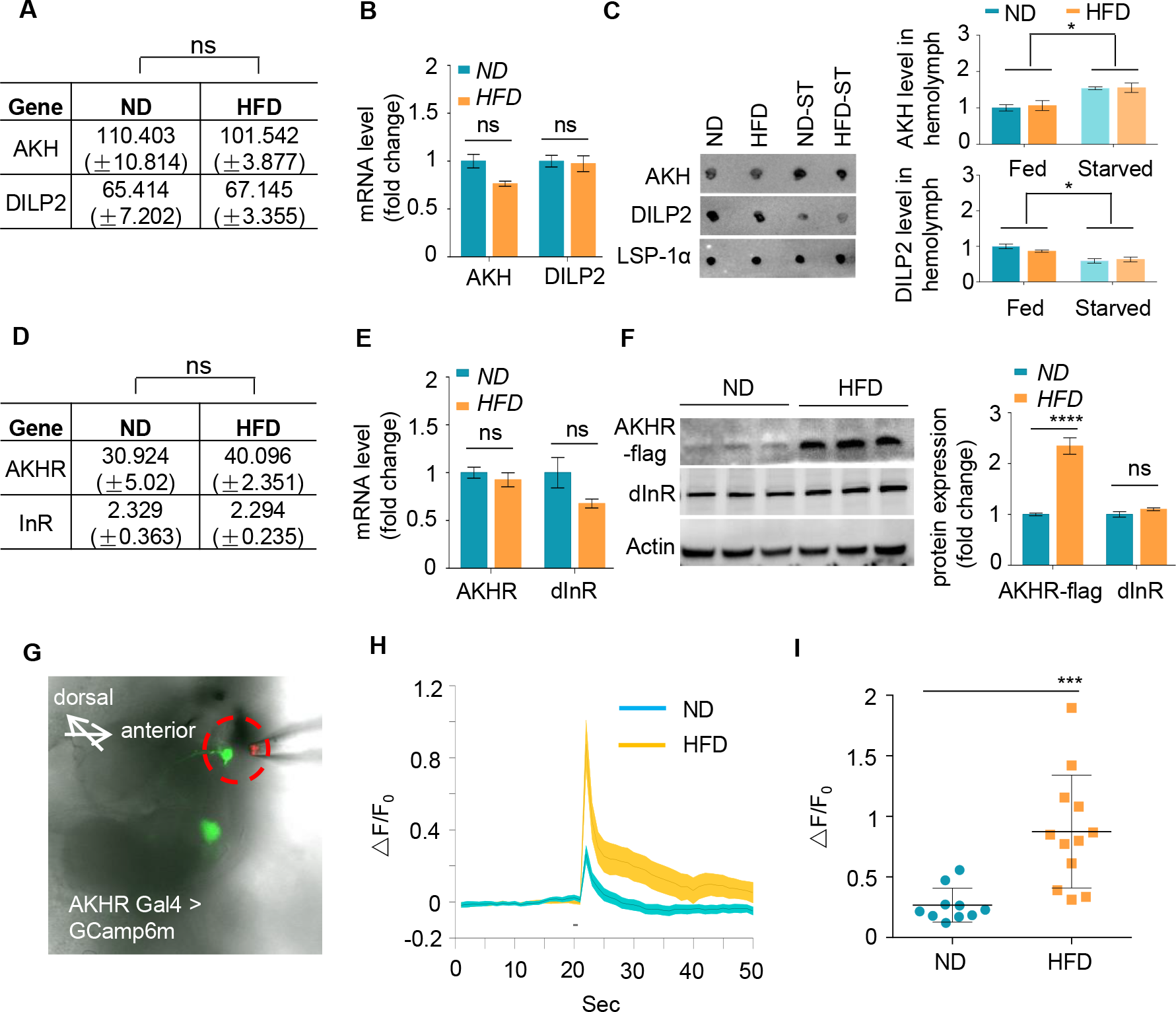
HFD increases neuronal AKHR protein. (A-B) AKH and DILP2 mRNA levels of wild-type virgin female flies fed with ND or HFD for 5 days. The fly heads and associated tissues were collected and subjected to RNAseq (A) or quantitative RT-PCR (B) (n = 3). (C) AKH and DILP2 protein levels in the hemolymph of wild-type virgin female flies, fed and starved (n = 3). Hemolymph samples were collected from the flies and AKH and DILP2 protein levels were analyzed with dot blot by using AKH and DILP2 antibodies. (D-E) AKHR and InR mRNA levels were analyzed similar to A-B (n = 3). (F) AKHR and dInR protein levels of fly heads were analyzed by western blot (n = 3). Note that AKHR-flag knock-in flies were generated by inserting FLAG sequence into the C terminal of AKHR gene via CRISPR/cas9 mediated gene editing. Antibodies against FLAG and dInR were used in the western blot. (G) Schematic diagram of the *ex vivo* calcium imaging. Green signals indicate AKHR^+^ neurons expressing GCaMP6 (*AKHR:BD/+; nSyb:AD/UAS-GCaMP6m*). Dotted red circle indicates the position of the pipette delivering synthetic AKH. (H-I) Representative traces (H) and quantification (I) of peak calcium transients of AKHR^+^ neurons upon AKH administration (n = 10-12). ns, p > 0.05; *p < 0.05; ***p < 0.001. Student’s t-test and one-way ANOVA followed by post hoc test weith Bonferroni correction were used for pair-wise and multiple comparisons, respectively.

We then asked whether the cognate receptors of AKH and DILP2, named AKHR and dInR (*Drosophila* insulin-like receptor), were affected by HFD feeding. While AKHR mRNA levels remained unchanged after HFD feeding, as revealed by RNAseq and quantitative RT-PCR (Figure 2D-E), AKHR protein was significantly elevated in the head tissue of HFD-fed flies vs. that of ND-fed flies, as demonstrated by western blot (Figure 2F). The production of dInR was not affected by HFD at both the transcriptional and translational levels (Figure 2D-F). These data suggest that HFD exerts a robust effect on AKHR protein accumulation without altering its mRNA production in the fly head.

We have previously shown that AKHR was expressed in two clusters of OA neurons in the subesophageal zone (SEZ) region of the fly brain (Yu et al., 2016). These neurons were responsive to the hunger hormone AKH and exerted a robust effect in inducing starvation-induced hyperactivity. Since AKHR protein levels were upregulated by HFD feeding, we reasoned that these AKHR^+^ neurons might be more sensitive to AKH upon HFD feeding. Indeed, by using a calcium imaging setup with *ex vivo* brain preparations, we found that these AKHR^+^ neurons in HFD-fed flies exhibited more robust calcium transients when stimulated with synthetic AKH than those in ND-fed flies (Figure 2G-I). Taken together, HFD leads to increased AKHR protein accumulation in the fly brain and enhanced sensitivity of AKHR^+^ neurons to the hunger hormone AKH. These results may account for the observations that HFD-fed flies become more sensitive to starvation and exhibit enhanced hyperactivity upon starvation (Figure 1).

### HFD enhanced AKHR accumulation by suppressing neuronal autophagy

We then sought to understand how HFD enhanced AKHR protein accumulation without promoting its mRNA production. One plausible hypothesis was that the degradation of AKHR via lysosome and/or proteasome was suppressed by HFD feeding. To examine this possibility, we ectopically expressed HA-tagged AKHR protein in cultured *Drosophila* S2 cells, and found that blocking lysosome-mediated protein degradation by lysosome inhibitors chloroquine (CQ) and NH_4_Cl increased AKHR protein levels dramatically, while MG132 treatment to disturb proteasome function exerted no effect (Figure 3A). Therefore, AKHR is specifically degraded via the lysosome pathway.

**Figure 3.**
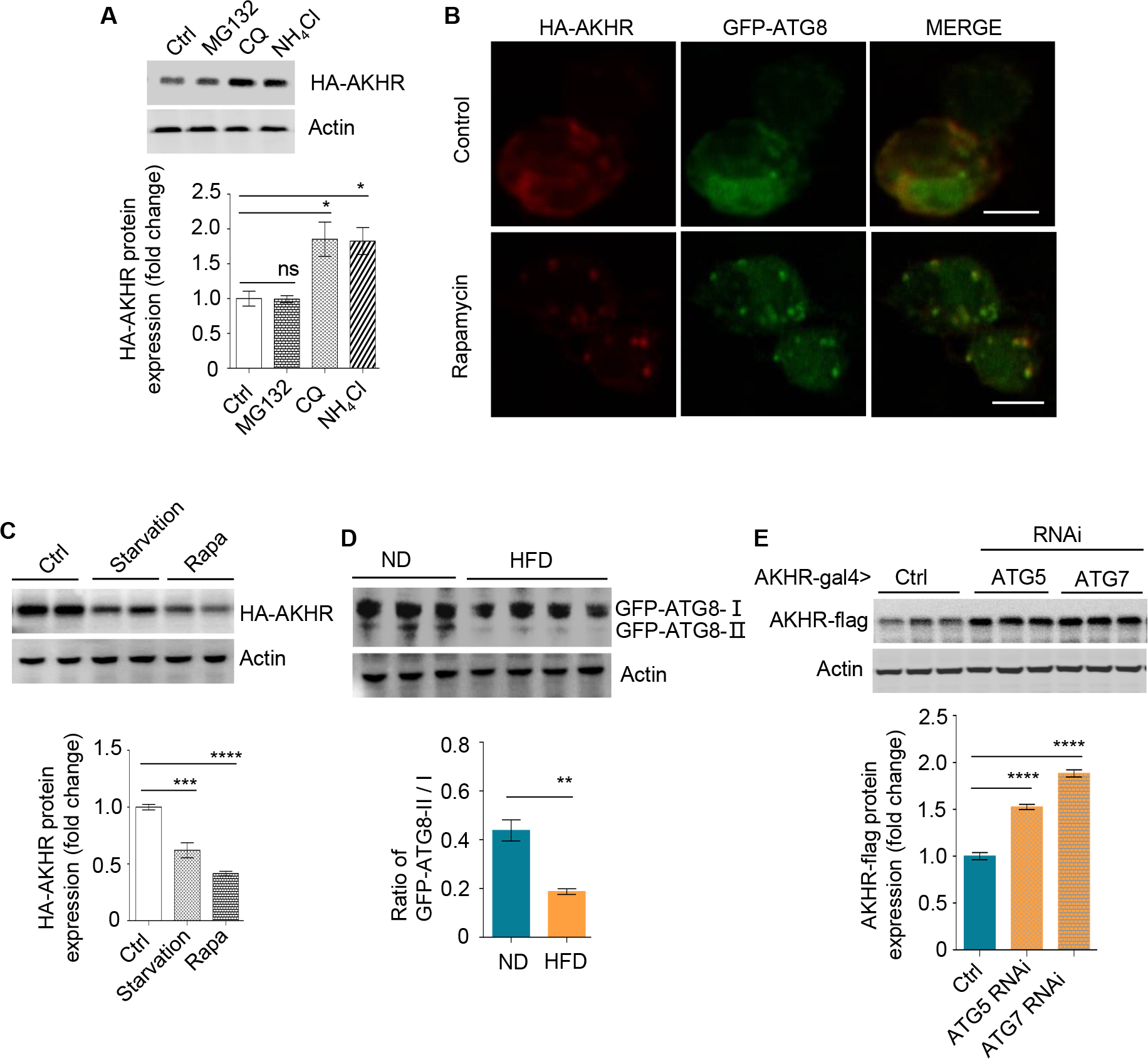
AKHR accumulation is induced by the suppression of autophagy. (A) Quantification of AKHR protein levels in cultured *Drosophila* S2 cells upon the treatment of proteasome and lysosome inhibitors (n = 3). Cultured cells transiently expressing AKHR-HA were treated with the indicated chemicals. The lysates were analyzed by western blot with HA antibody. (B) Formation of autophagosome in cultured S2 cells. Cultured cells transiently expressing AKHR-HA and GFP-ATG8 were treated with 200 nM rapamycin for 12 hr and stained for HA antibody. (C) Quantification of AKHR protein levels in Cultured *Drosophila* S2 cells upon starvation and the treatment of rapamycin (n = 3). Cultured S2 cells transiently expressing AKHR-HA were starved or treated with rapamycin, and AKHR-HA was analyzed by western blot. (D) Quantification of ATG8 activation in the head tissues of flies *in vivo* (n = 3-4). GFP-ATG8 was specifically expressed in AKHR^+^ neurons by using the GAL4-UAS system. Flies were fed with ND or HFD and their heads were collected for western blot with GFP antibody. The ratio between ATG8-II and ATG8-I bands indicates the level of autophagy. (E) Quantification of AKHR protein levels upon suppressing neuronal autophagy in AKHR^+^ neurons *in vivo* (n = 3). ATG5 and ATG7 were knocked down by the expression of RNAi constructs in AKHR^+^ neurons. AKHR-flag expression was assayed by western blot. ns, p >0.05; *p < 0.05; **p < 0.01; ****p < 0.0001. Student’s t-test and one-way ANOVA followed by post hoc test weith Bonferroni correction were used for pair-wise and multiple comparisons, respectively.

We next asked whether AKHR was degraded via autophagy, a specific lysosome-dependent protein degradation pathway (Klionsky, 2007). When treated with rapamycin, a robust inducer of autophagy, cultured S2 cells quickly formed characteristic autophagosomes, as indicated by the formation of dotted fluorescent GFP-ATG8 (the fly homolog of mammalian LC3) (Figure 3B, green). Notably, upon rapamycin treatment, HA-tagged AKHR protein was also enriched in these autophagosomes (Figure 3B, red). Moreover, inducing autophagy by both starvation and rapamycin resulted in significantly decreased AKHR protein levels in cultured S2 cells (Figure 3C). Therefore, the degradation of AKHR was solely mediated by the lysosome-dependent autophagy pathway *in vitro*. Notably, although it was widely accepted that membrane proteins were mainly degraded by endocytosis and then lysosome (e.g. LDLR and EGFR) (Beguinot et al., 1984), it has also been reported that certain membrane proteins can be degraded via autophagy, such as GABA and AMPA receptor subunits (Jin et al., 2018).

We then aimed to confirm these *in vitro* results and asked whether HFD induced AKHR accumulation in AKHR^+^ neurons via suppressing its degradation by the autophagy pathway *in vivo*. Indeed, HFD suppressed autophagy in the AKHR^+^ neurons, as evident by the decrease of membrane-associated ATG8 upon HFD feeding (lower band of GFP-ATG8, Figure 3D). Additionally, suppressing autophagy specifically in AKHR^+^ neurons, by RNAi knock-down of two critical autophagy initiators ATG5 and ATG7, significantly enhanced AKHR accumulation (Figure 3E), phenocopying the effect of HFD feeding. Taken together, HFD feeding suppresses neuronal autophagy and enhances AKHR accumulation in AKHR^+^ neurons.

### Suppression of autophagy in AKHR^+^ neurons enhanced their sensitivity to starvation

Since HFD enhanced AKHR accumulation in AKHR^+^ neurons via suppressing autophagy, we hypothesized that blocking autophagy in AKHR^+^ neurons in ND-fed flies would mimic the effect of HFD feeding, leading to higher AKHR accumulation, enhanced AKH sensitivity, and hence stronger starvation-induced hyperactivity.

We first tested this hypothesis in the calcium imaging setup using the *ex vivo* brain preparation. Indeed, RNAi knock-down of ATG5 and ATG7 in AKHR^+^ neurons led to significantly enhanced calcium transients upon AKH administration in ND-fed flies (Figure 4A), phenocopying the effect of HFD feeding.

**Figure 4.**
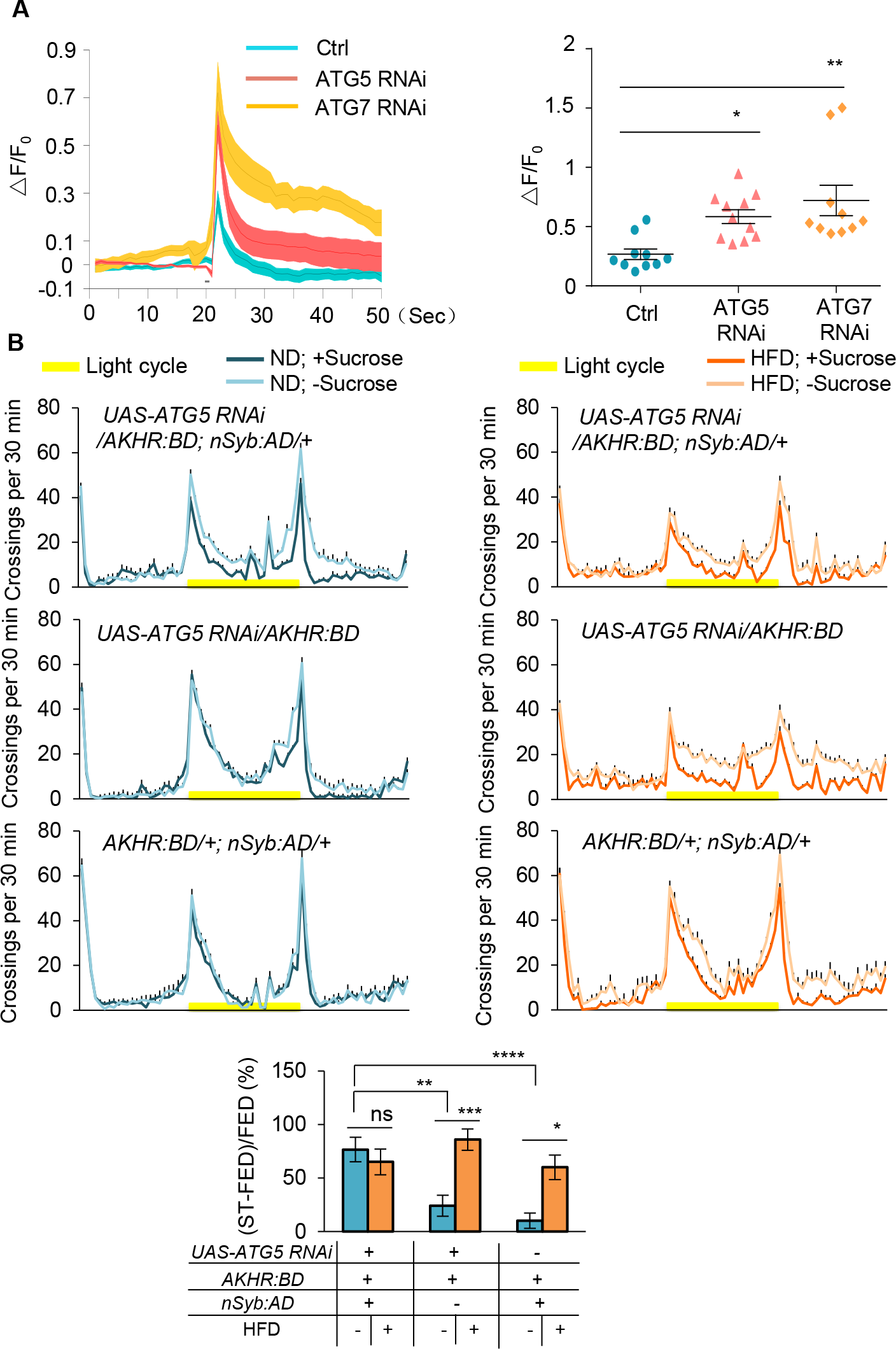
Inhibition of autophagy increases starvation-induced hyperactivity. (A) Representative traces (left) and quantification (right) of peak calcium responses of AKHR^+^ neurons to AKH upon suppressing neuronal autophagy in AKHR^+^ neurons (n = 10-11). The detailed genotypes: *UAS-ATG5(7) RNAi/AKHR:BD; nSyb:AD/UAS-GCaMP6m*. For the Ctrl group, *UAS-RNAi* transgene was not introduced. (B) Midline crossing activity (upper) of indicated genotypes assayed in the presence or absence of 5% sucrose in Day 1 (n = 43-62). Starvation-induced hyperactivity after 5-day HFD feeding vs. ND feeding experience is also shown (lower). ns, p > 0.05; *p < 0.05; **p < 0.01; ***p < 0.001; ****p < 0.0001. Student’s t-test and one-way ANOVA followed by post hoc test weith Bonferroni correction were used for pair-wise and multiple comparisons, respectively.

Consistent with these results, blocking autophagy in AKHR^+^ neurons via RNAi knock-down of ATG5 significantly enhanced starvation-induced hyperactivity in ND-fed flies (Figure 4B), again phenocopying the effect of HFD feeding. ATG7 knock-down exerted similar behavioral results (Figure 4-figure supplement 1). In both conditions, HFD feeding did not further enhance starvation-induced hyperactivity (Figure 4B and Figure 4-figure supplement 1). Taken together, suppressing neuronal autophagy in AKHR^+^ neurons resembles the effect of HFD feeding, enhancing the sensitivity of AKHR^+^ neurons as well as starvation-induced hyperactivity.

### HFD suppressed autophagy via activating AMPK-TOR signaling

To further examine the cellular mechanism underlying HFD-induced suppression of neuronal autophagy, we carried out RNAseq analysis of fly heads harvested from ND-fed and HFD-fed flies. As shown in Figure 5-figure supplement 1 and 2, we found that a collection of genes related to anabolic activities were upregulated while a collection of catabolic genes were downregulated in HFD-fed flies. These results hinted a possibility that TOR signaling, the key regulator of anabolism, was activated by HFD. Notably, TOR signaling is also a robust inducer of autophagy (Wullschleger et al., 2006). Therefore, TOR signaling may mediate the effect of HFD on autophagy.

**Figure 5.**
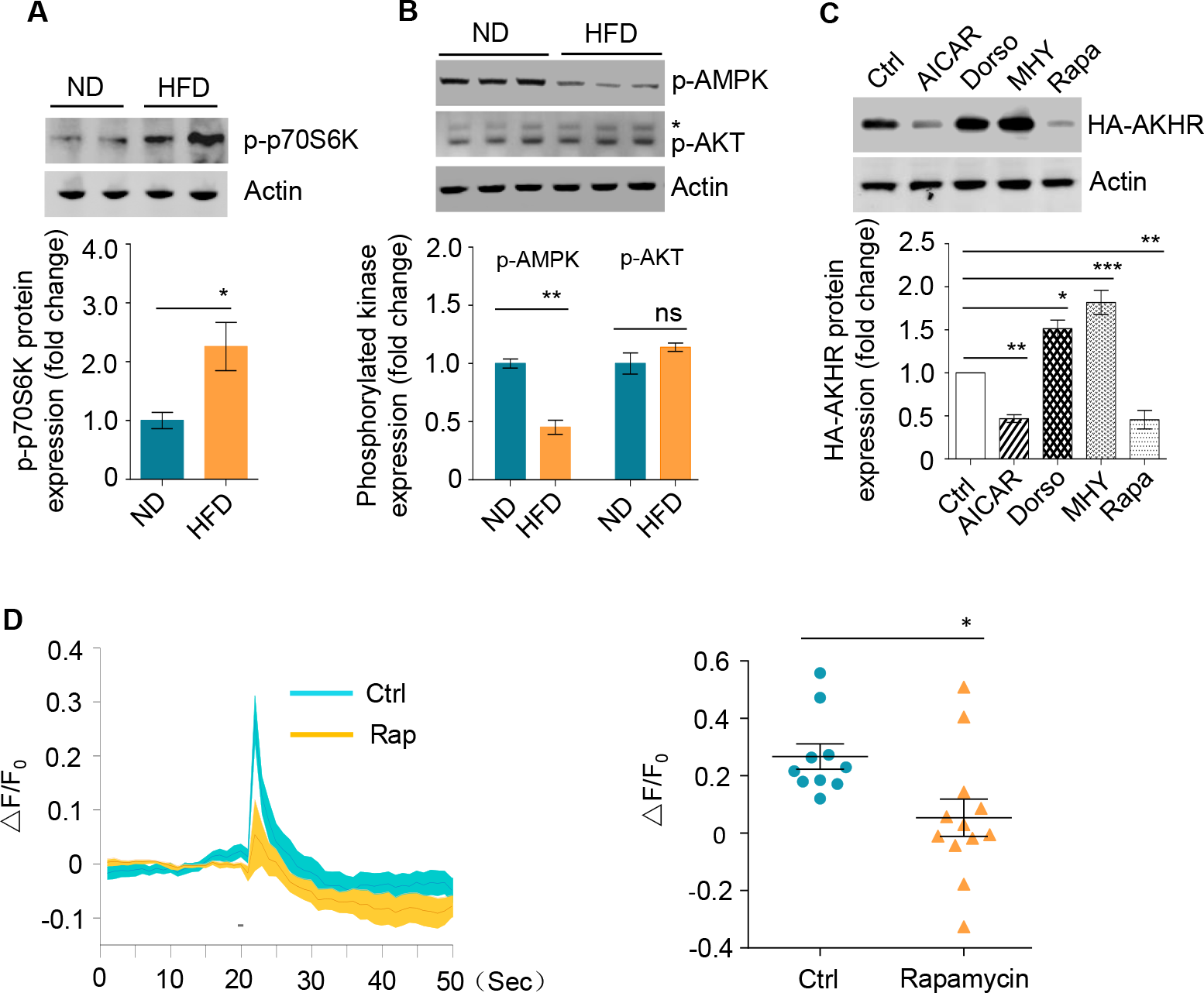
HFD activates TOR signaling. (A-B) p70s6k/AKT/AMPK phosphorylation under ND vs. HFD feeding conditions *in vivo*. The head tissues of wild-type flies fed with ND or HFD were harvested and subjected to western blot with phosphorylated p70s6k antibody (A) and AKT/AMPK antibodies (B) (n = 3). *Nonspecific bands. (C) AKHR accumulation upon pharmacological manipulation of AMPK-TOR signaling in cultured *Drosophila* S2 cells (n = 3). Cultured S2 cells transiently expressing AKHR-HA were treated with the indicated chemicals and then analyzed by western blot with HA antibody. (D) Representative traces (left) and quantification (right) of peak calcium responses of AKHR^+^ neurons to AKH, with or without rapamycin treatment (n = 10-12). ns, p > 0.05; *p < 0.05; **p < 0.01; ***p < 0.001. Student’s t-test and one-way ANOVA followed by post hoc test weith Bonferroni correction were used for pair-wise and multiple comparisons, respectively.

To further confirm this hypothesis, we examined p70S6K phosphorylation *in vivo*, which was a reliable indicator of TOR activation (Wullschleger et al., 2006). Indeed, the head tissues of HFD-fed flies exhibited higher levels of p70S6K phosphorylation (Figure 5A), suggesting that HFD indeed activated TOR signaling. TOR signaling could be modulated by two major upstream signals, negatively by AMPK (AMP-activated protein kinase) signaling and positively by AKT signaling (Wullschleger et al., 2006). We found that upon HFD feeding, AMPK signaling was suppressed, whereas AKT signaling remained unaffected (Figure 5B), suggesting that HFD modulated TOR signaling via the suppression of AMPK signaling.

Since AMPK-TOR signaling was known to modulate autophagy and the degradation of specific proteins, it was therefore possible that HFD suppressed neuronal autophagy and hence enhanced AKHR accumulation via modulating AMPK-TOR signaling. If this was the case, manipulating AMPK-TOR signaling would exert robust effect on AKHR accumulation just as HFD-feeding. Indeed, pharmacological activation of TOR signaling by MHY and suppression of AMPK by Dorsomorphin both resulted in AKHR accumulation in cultured S2 cells, whereas suppression of TOR signaling by rapamycin and activation of AMPK by AICAR exerted the opposite effect (Figure 5C). Consistent with these *in vitro* results, rapamycin treatment in the *ex vivo* brain preparation led to decreased AKH sensitivity in AKHR^+^ neurons (Figure 5D), further confirming that AMPK-TOR signaling plays an important role in regulating AKHR accumulation and hence the sensitivity of AKHR^+^ neurons.

### AMPK-TOR signaling mediated the effect of HFD on starvation-induced hyperactivity

Since AMPK-TOR signaling mediated the effect of HFD on AKHR accumulation, we reasoned that manipulating AMPK-TOR signaling *in vivo* would affect the onset of starvation-induced hyperactivity.

Indeed, suppressing TOR signaling in AKHR^+^ neurons by ectopic expression of dominant negative TOR^TED^ (Hennig and Neufeld, 2002) resulted in significantly decreased starvation-induced hyperactivity in ND-fed flies, and HFD feeding did not further enhance starvation-induced hyperactivity in these flies (Figure 6A). Rapamycin feeding exerted similar effect (Figure 6B). These results were consistent with our findings that suppression of TOR signaling led to much reduced AKHR levels (Figure 5C) and AKH sensitivity (Figure 5D).

**Figure 6.**
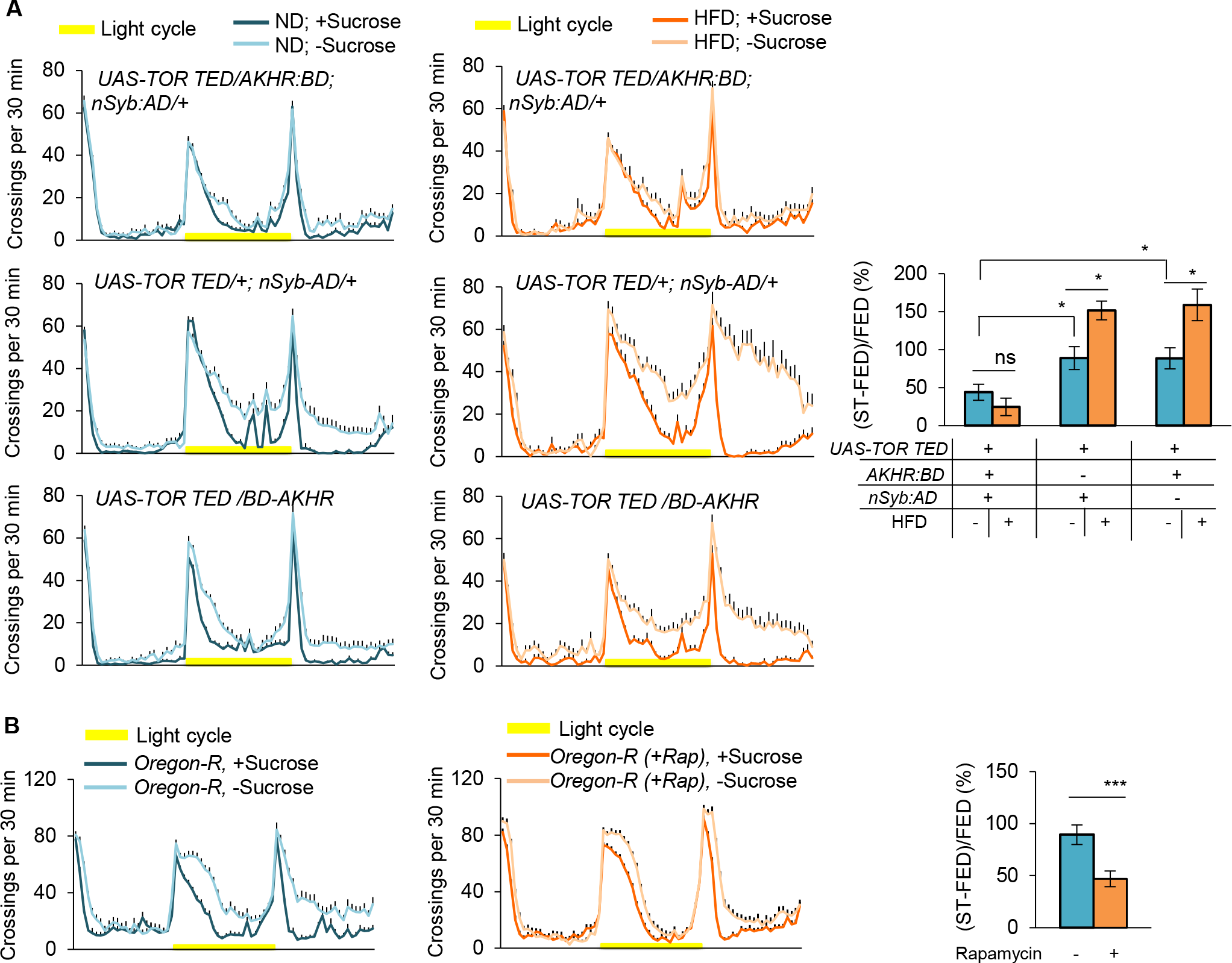
AMPK-TOR signaling modulates starvation-induced hyperactivity. (A) Midline crossing activity of indicated genotypes assayed in the presence or absence of 5% sucrose in Day 1 (left) (n = 35-58). Starvation-induced hyperactivity after 5-day HFD feeding vs. ND feeding experience is also shown (right). (B) Midline crossing activity of wild-type virgin females (pre-fed with or without rapamycin) assayed in the presence or absence of 5% sucrose in Day 1 (left) (n = 48-64). Starvation-induced hyperactivity after 5-day HFD feeding vs. ND feeding experience is also shown (right). ns, p > 0.05; *p < 0.05; **p < 0.01; ***p < 0.001. Student’s t-test and one-way ANOVA followed by post hoc test weith Bonferroni correction were used for pair-wise and multiple comparisons, respectively.

To further confirm these results, we genetically activated TOR signaling in AKHR^+^ neurons, by RNAi knock-down of TSC1, a suppressor of TOR signaling (Wullschleger et al., 2006) and by the ectopic expression of a dominant negative form of AMPK protein (AMPK^DN^) (Wullschleger et al., 2006). Both manipulations resulted in upregulated starvation-induced hyperactivity in ND-fed flies (Figure 6-figure supplement 1 and 2), phenocopying the effect of HFD feeding (Figure 1A). Collectively, these data further confirm that HFD enhances starvation-induced hyperactivity via modulating AMPK-TOR signaling in AKHR^+^ neurons.

### A lipoprotein LTP and its cognate receptor LpR1 linked HFD feeding to enhanced starvation-induced hyperactivity

We next sought to understand how HFD feeding exerted a robust effect on the autophagy pathway of a small group of brain neurons expressing AKHR. Since dietary lipids needed to enter the circulating system and different fly organs via specific carriers named lipoproteins, we asked whether certain lipoprotein(s) played a role in mediating the behavioral effect of HFD feeding.

Three lipophorins, the main *Drosophila* lipoprotein, circulate in the hemolymph and transport lipids to different fly organs (Rodriguez-Vazquez et al., 2015). We performed proteomic analysis of circulating proteins in flies’ hemolymph by mass spectrometry and found that only one of the three lipoproteins, named Lipid Transfer Particle (LTP), but not the other two [Lipophorin (LPP) and Microsomal Triglyceride Transfer Protein (MTP)], was significantly upregulated in HFD-fed flies vs. ND-fed flies (Figure 7A-7B). Meanwhile, both triglycerides and cholesterol were enriched in the hemolymph of HFD-fed flies (Figure 7C-7D). Therefore, it was possible that LTP was the major carrier for excess circulating lipids after HFD feeding.

**Figure 7.**
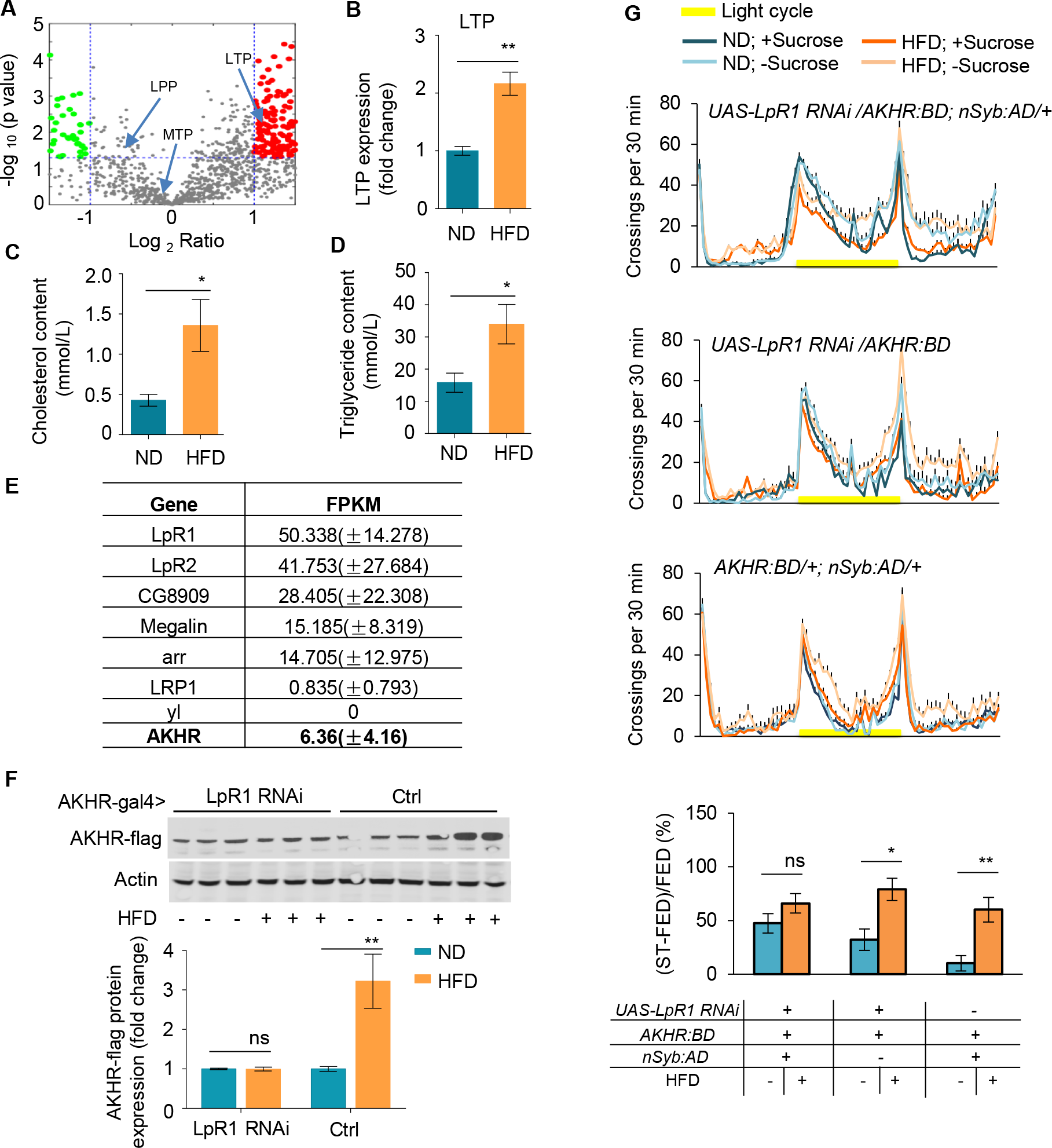
LpR1 is required for HFD-strengthened hyperactivity under starvation. (A-B) LTP protein levels in the hemolymph. Hemolymph was collected from flies fed with ND or HFD and subjected to LC-MS/MS analysis. Volcano plot (A) shows peptides with differential enrichment (under ND vs. HFD feeding conditions). The horizontal line indicates the significance threshold (p = 0.01) and the vertical lines indicate two-fold change. Green and red dots represent down-regulated and up-regulated peptides, respectively. Arrows indicate three lipoproteins (LTP, LPP and MTP). The level of LTP was shown in B (n = 3). (C-D) The cholesterol level (C) and triglyceride level (D) in the hemolymph harvested from ND vs. HFD fed flies (n= 4-6). (E) Single cell RNAseq shows the gene expression level of candidate lipoprotein receptors in AKHR^+^ neurons (n = 5). (F) AKHR protein levels from the head tissues of the indicated genotypes fed with ND or HFD (n = 3). (G) Midline crossing activity of indicated genotypes assayed in the presence or absence of 5% sucrose in Day 1 (upper) (n = 43-57). Starvation-induced hyperactivity after 5-day HFD feeding vs. ND feeding experience is also shown (lower). ns, p > 0.05; *p < 0.05; **p < 0.01. Student’s t-test and one-way ANOVA followed by post hoc test weith Bonferroni correction were used for pair-wise and multiple comparisons, respectively.

Recruitment of lipoproteins to cell membranes mediated by its receptors is a key event that initiates the transfer of neutral lipids to cells (Rodriguez-Vazquez et al., 2015). To understand the role of excess circulating lipids in suppressing autophagy of AKHR^+^ neurons, it was necessary to identify lipoprotein receptor expressed in these cells. To this aim, we carried out single cell RNAseq analysis for individual AKHR^+^ neurons harvested from the fly brains *in situ*. Among seven identified lipophorin receptors in *Drosophila*, lipophorin receptor 1 (LpR1) was most abundantly expressed in AKHR^+^ neurons (Figure 7E).

We thus examined the potential role of LpR1 in mediating the effect of HFD feeding on AKHR accumulation and starvation-induced hyperactivity. RNAi knock-down of LpR1 in AKHR^+^ neurons eliminated the effect of HFD on AKHR accumulation (Figure 7F). Consistently, these flies did not show an enhancement of starvation-induced hyperactivity by HFD feeding (Figure 7G). Therefore, we concluded that a specific lipoprotein LTP and its cognate receptor LpR1 mediated the effect of HFD on neuronal autophagy, AKHR protein accumulation, and therefore the onset of starvation-induced hyperactivity upon starvation.

### Diet-independent hyperlipidemia also enhanced starvation-induced hyperactivity

HFD is known to be a robust reward signal and can induce food craving (de Macedo et al., 2016). We therefore asked whether enhanced starvation-induced hyperactivity by HFD feeding was solely mediated by excess lipid uptake, or HFD feeding also exerted other non-metabolic effects. By knocking down LpR1 in the fat body (*ppl-GAL4/UAS-LpR1 RNAi*), the major organ for lipid storage and metabolism, we generated diet-independent hyperlipidemia in ND-fed flies (Figure 8A-B). Notably, similar effect was also observed in mouse models (Wang et al., 2016)

**Figure 8.**
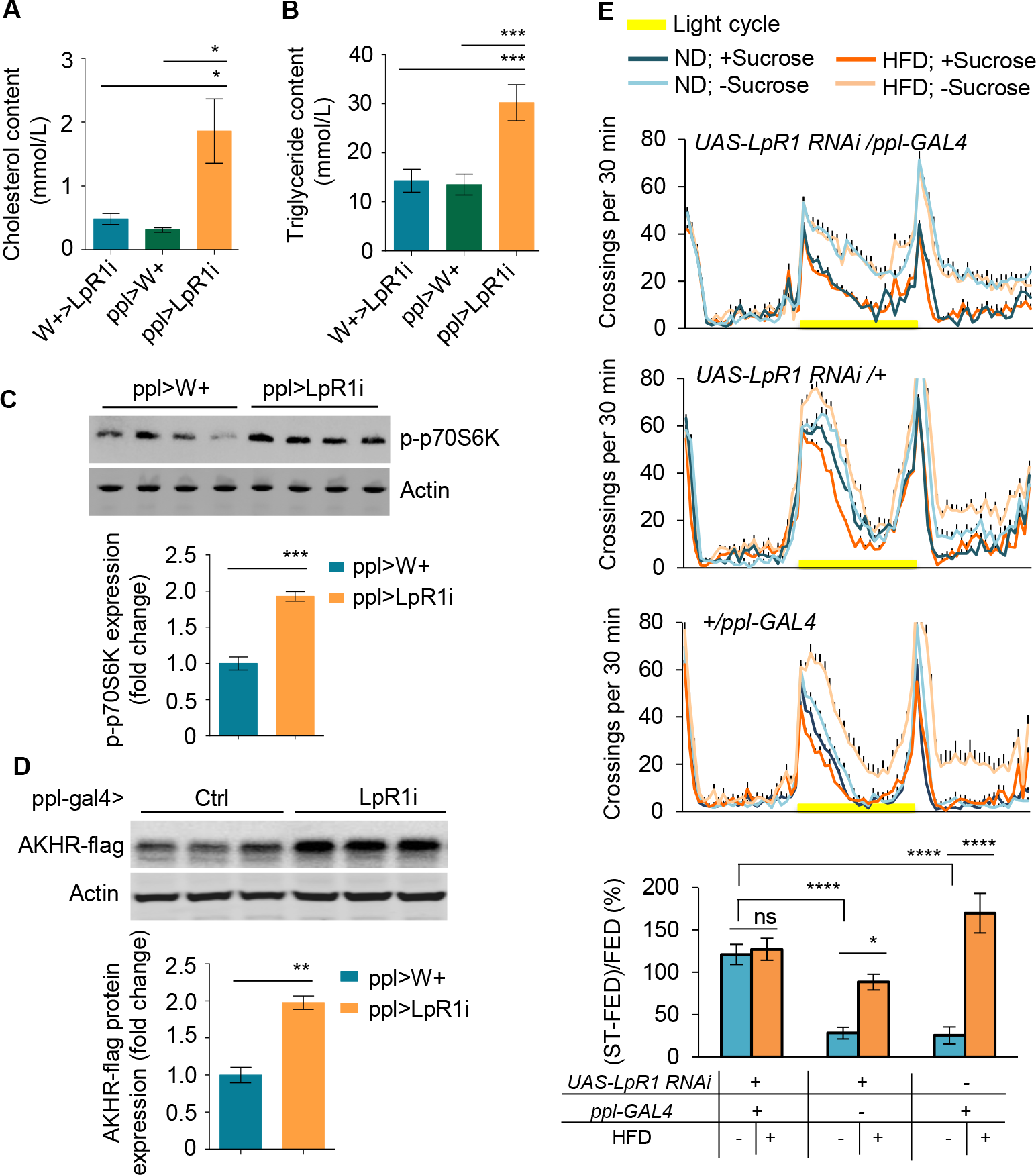
Diet-independent hyperlipidemia promotes the excitability of AKHR^+^ neurons and starvation-induced hyperactivity. (A-B) Average levels of cholesterol (A) and triglyceride (B) in the hemolymph of the indicated genotypes (n = 4-11). (C-D) The levels of phosphorylated p70s6k (C) and AKHR-flag (D) from the head tissues of indicated genotypes (n = 4). (E) Midline crossing activity of indicated genotypes assayed in the presence or absence of 5% sucrose in Day 1 (upper) (n = 41-63). Starvation-induced hyperactivity after 5-day HFD feeding vs. ND feeding experience is also shown (lower). ns, p > 0.05; *p < 0.05; ***p < 0.001; ****p < 0.0001. Student’s t-test and one-way ANOVA followed by post hoc test weith Bonferroni correction were used for pair-wise and multiple comparisons, respectively.

We then asked whether diet-independent hyperlipidemia had similar effects on AKHR accumulation and starvation-induced hyperactivity as HFD feeding. Compared with the control group, the phosphorylation of p70S6K in the diet-independent hyperlipidemia group was significantly up-regulated (Figure 8C). These results showed that like HFD feeding, diet-independent hyperlipidemia also activated TOR signaling (Figure 8C). In addition, diet-independent hyperlipidemia led to upregulated AKHR accumulation (Figure 8D), mimicking the effect of HFD feeding. Consistent with these results, these ND-fed, hyperlipidemic flies showed significantly stronger starvation-induced hyperactivity than the two control groups (Figure 8E), which was comparable to HFD-fed flies (Figure 1A). Taken together, diet-independent hyperlipidemia phenocopies the effect of HFD feeding, further confirming that HFD feeding reshapes the circuitry underlying starvation-induced hyperactivity through excess dietary lipid intake.

## DISCUSSION

There is accumulating evidence that notes the effect of HFD on food consumption from insects to human, which results in obesity and obesity-associated metabolic diseases. But the effect of HFD on another critical food intake related behavior, food seeking, remains largely uncharacterized. Conceptually, food-seeking behavior in the fruit fly is composed of two behavioral components, increased sensitivity to food cues, and enhanced exploratory locomotion, which altogether facilitates the localization and acquisition of desirable food sources (Root et al., 2011; Yu et al., 2016). Previous work from our lab has shown that starvation promotes starvation-induced hyperactivity, the exploratory component of food-seeking behavior, in fruit flies via a small group of OA neurons in the fly brain. These hunger-sensing OA neurons sample the metabolic status by detecting two groups of functionally antagonistic hormones, AKH and DILPs, and promote starvation-induced hyperactivity (Yu et al., 2016).

In this present study, we demonstrated that this behavior was affected by metabolic challenges. After a few days of HFD feeding, flies became behaviorally hypersensitive to starvation and as a result their starvation-induced hyperactivity was greatly enhanced, despite that their energy storage was not reduced and their food consumption was not affected. These results suggested that HFD feeding might specifically modulate the activity of the neural circuitry underlying starvation-induced hyperactivity and offered us an opportunity to further elucidate the cellular and circuitry mechanisms underlying behavioral abnormalities upon metabolic challenges.

As an insect counterpart of mammalian glucagon, AKH acts as a hunger signal to activate its cognate receptor AKHR expressed in the fat body and subsequently triggers lipid mobilization and energy allocation (Bharucha et al., 2008). In the fly brain, a small number of OA neurons also express AKHR. We and other labs have shown that these neurons are responsive to starvation and modulate various behaviors including food seeking and drinking (Jourjine et al., 2016; Yu et al., 2016). In that sense, these OA^+^AKHR^+^ neurons are functionally analogous to mammalian NPY/AgRP neurons in the hypothalamus, which also senses organismal metabolic states and regulates specific food intake behaviors. In this present study, we found that OA^+^AKHR^+^ neurons exhibited a higher AKHR protein accumulation and became hypersensitive to AKH after HFD feeding. Notably, HFD feeding in mammals also increases the excitability of NPY/AgRP neurons, which contributes to the hypersensitivity to starvation and increased food consumption (Vernia et al., 2016). Thus, HFD may play a conserved role in the regulation of neuronal excitability and food intake related behaviors in both fruit flies and mammals.

Autophagy, a lysosomal degradative process that maintains cellular homeostasis, is critical for energy homeostasis. Upon cellular starvation, autophagy generates additional energy supply by breaking down macromolecules and subcellular organelles (Klionsky, 2007). At the organismal level, autophagy also contributes to the regulation of food intake and hence organismal energy homeostasis. For example, fasting induces autophagy in NPY/AgRP neurons via fatty acid uptake and promotes AgRP expression, which in turn enhances food intake (Kaushik et al., 2011). In line with these results, eliminating autophagy in NPY/AgRP neurons reduces food intake and hence body weight and fat deposits (Kaushik et al., 2011). Conversely, loss of autophagy in POMC neurons displays increased food intake and adiposity (Coupe et al., 2012). In this present study, we showed that in fruit flies neuronal autophagy is critical for the function of OA^+^AKHR^+^ neurons to sense hunger and regulate starvation-induced hyperactivity. Meanwhile, HFD feeding attenuated autophagy in these OA^+^AKHR^+^ neurons and exerted behavioral and metabolic effects.

Accumulating evidence suggests that HFD suppresses autophagy in different peripheral tissue types such as liver, skeletal muscle, and the adipose tissue (Feng et al., 2016; He et al., 2012; Liu et al., 2009). Similarly, HFD suppresses autophagy in the hypothalamus, whereas blocking hypothalamic autophagy, particularly in POMC neurons, exacerbates HFD induced obesity (Coupe et al., 2012). In this present study, we showed that HFD suppressed neuronal autophagy in OA^+^AKHR^+^ neurons and enhanced AKHR accumulation in these neurons. As a result, OA^+^AKHR^+^ neurons became hypersensitive to starvation and enhanced starvation-induced hyperactivity. It would be of interest to examine whether HFD also reduces autophagy and increases the accumulation of specific membrane receptors in mammalian NPY/AgRP neurons.

We also sought to examine the cellular mechanism that linked HFD feeding to the reduction of autophagy. We found that HFD feeding activated TOR signaling as evident by both RNAseq and western blot. TOR, a highly conserved serine-threonine kinase, controls numerous anabolic cellular processes. We found in this present study that TOR signaling was tightly associated with the activity of AKHR^+^ neurons and the behavioral responses upon HFD feeding. Genetic enhancement of TOR activity in AKHR^+^ neurons increased AKHR protein accumulation, the sensitivity of these neurons to AKH, and hence starvation-induced hyperactivity, all of which mimicked the effect of HFD feeding. Inhibiting TOR activity exerted an opposite effect. In addition, we also found that the effect of HFD on TOR signaling was mediated by AMPK signaling. These results altogether suggest that AMPK-TOR signaling in AKHR^+^ neurons plays an important role in maintaining the homeostasis of these neurons and determining the responsiveness to HFD feeding. Similarly, rodent studies have shown that manipulating AMPK-TOR signaling resulted in the dysfunction of NPY/AgRP neurons as well as POMC neurons, which led to abnormal food consumption and adiposity (Kong et al., 2016; Yang et al., 2012). It would be of interest to examine whether HFD also modulates AMPK-TOR signaling in specific hypothalamic neurons.

We next sought to understand how AKHR^+^ neurons detected HFD, or more specifically, excess lipid ingested by the flies. As an essential nutrient and important energy reserve, dietary lipids were transported via their carrier proteins, named lipoproteins, in the circulation system and regulate multiple cellular signaling pathways. Proteomic analysis helped us to identify that one lipoprotein LTP was enriched in flies’ hemolymph after HFD feeding. Single-cell RNAseq of AKHR^+^ neurons identified a number of lipoprotein receptors, especially LpR1, were highly expressed in these neurons. Therefore, we proposed that AKHR^+^ neurons might sense HFD feeding via LTP-LpR1 signaling. Evidently, we found that eliminating LpR1 in AKHR^+^ neurons could protect flies from HFD, reducing AKHR accumulation and starvation-induced hyperactivity. Conversely, eliminating LpR1 in the fat body, the major lipid reservoir of flies, created diet-independent hyperlipidemia and mimicked the effect of HFD feeding on flies’ starvation-induced hyperactivity. Taken together, we propose a working model that upon HFD feeding, excess dietary lipids are transported by LTP in the hemolymph, which interacts with its cognate receptor LpR1 in OA^+^AKHR^+^ neurons. As a result, these neurons undergo a number of cellular signaling processes and eventually become hypersensitive to starvation (Figure 8-figure supplement 1).

To summarize, our present study established a link between an unhealthy diet and abnormalities of food intake related behaviors in a model organism. We have also deciphered the underlying mechanism involving intracellular AMPK-TOR signaling, reduced neuronal autophagy, accumulation of a specific hormone receptor, and increased excitability of a small group of hunger-sensing neurons. Our study will shed crucial light on the pathological changes in the central nervous system upon metabolic challenges. Given that the central control of metabolism and food intake related behaviors are highly conserved across different species, it will be of importance to further examine whether similar mechanisms also mediate the effect of HFD feeding on food intake and metabolic diseases in mammals.

## MATERIALS AND METHODS

### Flies

Flies were reared on standard fly food made of yeast, corn, and agar at 25°C and 60% humidity on a 12-hr light-12-hr dark cycle. Unless otherwise specified, virgin female flies were collected shortly after eclosion and placed in standard fly food (ND) vials (20 flies per vial) for 5-6 d prior to experiments. High fat diet (HFD)-fed flies were kept in vials with HFD medium containing 20% coconut oil and 80% ND for 5 d. The flies of rapamycin group were fed in ND containing 400 μM rapamycin for 5 d.

*nSyb-AD-AKHR-BD* was described previously (Yu et al., 2016). *UAS-TSC1 RNAi* (#5074), *UAS-ATG7 RNAi* (#2793) and *UAS-LpR1 RNAi* (#2568) were obtained from the Tsinghua Fly Center. *UAS-TOR^TED^* (#7013), *UAS-AMPK-DN* (#32112), *UAS-mCD8GFP* (#32186) and *ppl-GAL4* (#58768) were provided by the Bloomington *Drosophila* Stock Center at Indiana University. *UAS-ATG5 RNAi* flies were from Chao Tong (Zhejiang University).

### Chemicals, plasmids, and antibodies

If not otherwise indicated, all chemicals were from Sigma. MG132 (10 μM; Beyotime Biotechnology) and CQ (100 μM) exposure was for 4 hr. Rapamycin (200 nM; Sangon Biotech), AICAR (500 μM; Beyotime Biotechnology), MHY (5 μM) and Dorsomorphin (5 μM) were add to the medium for 12 hr.

pAC5.1-flag and pAC5.1-HA vectors were kindly provided by Xiaohang Yang (Zhejiang University). AKHR-HA was made by cloning the cDNA of AKHR into a pAC5.1-HA vector using SalI and NotI. The cDNA of ATG8 was from Chao Tong (Zhejiang University) and cloned into a pAC5.1-HA vector using EcoRI and BamHI to generate HA-ATG8.

The following antibodies were used: rabbit polyclonal antibodies to p-AKT, p-AMPK, HA, Flag, GFP and mouse monoclonal antibody to HA (Cell Signaling Technology); mouse monoclonal antibodies to Flag (MBL) and to actin (sigma); rabbit polyclonal antibody to AKH (biorbyt); rabbit polyclonal antibody to Dilp2 was kindly provided by Zhefeng Gong (Zhejiang University).

### Cell culture and transfection

S2 cells were grown in Schneider’s Insect Medium (sigma) supplemented with 10% fetal bovine serum (FBS) at 25°C. Transfections were performed using Lipofectamine 3000 according to the manufacturer’s instructions (Invitrogen). Cells were analyzed 24–48 hr after transfection.

### Behavioral assays

DAMS-based locomotion assay was performed as described in our earlier report (Yu et al., 2016). Briefly, individual virgin female flies were introduced into 5 mm × 65 mm polycarbonate tubes (Trikinetics) after being lightly anesthetized. One end of these tubes was filled with medium containing 2% (wt/vol) agar ± 5% (wt/vol) sucrose and the other end was blocked using cotton wool. These tubes were then inserted into DAMS monitors placing in fly incubators. The passage of flies through the middle of the tube was counted by an infrared beam through the midline of the tubes.

The MAFE assay was performed as described previously (Qi et al., 2015). Briefly, individual flies were gently aspirated and introduced into a 200 μL pipette tip. After exposing the proboscis by cutting the pipette tip, the flies were first sated with water and then presented with 100mM sucrose containing 2% blue dye (McCormick) filled in a graduated capillary (VWR, #53432-604). Until the flies became unresponsive to 10 serial food stimuli, food consumption was calculated according to the total volume of ingested food.

The FLIC assay was performed as described previously (Qi et al., 2015). Briefly, both feeding channels in the *Drosophila* Feeding Monitors (DFM) were filled with 5% sucrose. Individual flies were then gently aspirated into each feeding arena and their feeding activity was recorded for 24 hr. An electrical current generated by the physical contact between flies’ proboscis and the liquid food was recorded by the DFMs. From the original report, electrical current more than 120 a.u. was considered as actual feeding, and the total feeding bouts and the duration of feeding time were calculated accordingly.

### In vivo two-photon calcium imaging

The flies were lightly anesthetized on ice, and brains were dissected in oxygen saturation (95% O_2_, 5% CO_2_) buffer extracellular saline solution [103 mM NaCl, 3 mM KCl, 5 mM N-Tris (hydroxymethyl) methyl-2-aminoethane-sulfonic acid, 10 mM trehalose, 8 mM glucose, 26 mM NaHCO_3_, 1 mM NaH_2_PO_4_, 1.5 mM CaCl_2_, and 4 mM MgCl_2_, The pH was adjusted to 7.5 and osmolarity to 265 mOsm].

Dissected brains were adhered to a dish. Dorsal view is upward. Micropipettes (glass capillaries, Harvard Apparatus, 300092) with an opening diameter of approximately 10 μm were connected to a picospritzer III (Intracell). The tip of the pipette was positioned 50 μm to the cell body (Figure 2G). Air pressure on the picospritzer (III) was set to 3.3 psi, duration was 300 ms, the proper parameter required to eject solution from the pipette to induce the calcium response in AKHR promoter driven calcium indicator GCamp6m. Electrode was perfused by microloader. Alexa fluor 568 was mixed in the AKH solution (3.16 mM) to verify the eject area could cover the target AKHR neuron’s cell body (Figure 2G).

Images were acquired on Nikon’s Eclipse FN1 confocal microscopes with 40× /0.8 objectives. Time series were acquired at 1 Hz with 256 × 256 pixels and 12 bit, a resolution of 0.83 μm/ pixel, controlled by NIS elements. Laser power density on the scanned area of laser wavelength 488 nm was lower than 0.3 mW/ mm^2^, and 561 nm was 10 mW/mm^2^.

Total image acquired duration is 4 minute, first 1minute set to pre-puff drug time followed by 300 ms drug delivered. Image stacks were subsequently analyzed by Fiji (http://fiji.sc). ROIs were manually drawn around AKHR^+^ neuron cell body, and the intensity of 30 frames before puff time average to be F0, individual ΔF/F0 response traces were extracted, and the peak ΔF/F0 was obtained for significance analysis.

### Immunofluorescence staining

For immunostaining, Drosophila S2 cells were fixed in 4% formaldehyde. After washing thrice in PBS, cells were blocked in PBS/FBS (PBS, pH 7.4, containing 10% FBS). The cells were then incubated with appropriate primary and secondary antibodies in PBS/FBS with 0.1% saponin.

### RNAseq

Total RNA from fly heads was extracted from 5-day-old female flies using the Trizol reagent (Invitrogen, USA), subjected to poly(A) mRNA isolation, cDNA synthesization, library preparation (NEBNext Ultra DNA Library Prep Kit for Illumina, NEB), and sequencing (Illumina Hiseq2500/4000 platform). Sequence data were subsequently mapped to Drosophila genome and uniquely mapped reads were collected for further analysis. Gene expression was calculated by the RPKM (Reads Per Kilobase of exon per Million reads mapped). The genes with p-value less 0.05 were considered as the differentially expressed gene. The RNAseq data of fly heads were deposited in GEO under the accession code GSE129602.

### Single-cell RNAseq

As described in our previously report (Yu et al., 2016), individual AKHR^+^ cells (marked by GFP expression) were picked from dissected fly brain under a fluorescence microscope using a glass micropipette pulled from thick-walled borosilicate capillaries (BF120-69-10, Sutter Instruments). Separated cells were transferred to lysate buffer immediately, followed by reverse transcription and cDNA amplification (SMARTer Ultra Low RNA Kit for Sequencing, Clonetech). The amplified cDNA underwent library preparation using the NEBNext Ultra II DNALibrar kit and subject to sequencing by Illumina Hiseq 2500 platform. The sequenced raw data were purified to remove low-quality reads, adaptor sequences and amplification primer before mapping to *Drosophila* genome. Only mapped reads were selected for further analysis. FPKM (Fragments Per Kilobase Of Exon Per Million Fragments Mapped) was used to quantify gene expression. The single-cell RNAseq data were deposited in GEO under the accession code GSE129601.

### Measurements of CO2 production

The CO2 production was performed as described previously (Takeuchi et al., 2009). 1 ml plastic syringes were used for the respirometers after filling a small amount of CO2 absorbent (Soda lime, Sigma) between two pieces of sponge in the body of the syringe. A 5 μl fine graduated capillary connected to the plastic adaptor attached to the syringe. Five flies were gently aspirated into the syringe body and plunger inserted to form a closed chamber. After being kept on the flat surface at 25ºC for 15 min to equilibrate, a little ink was placed at the end of the micropipet. The CO2 production was calculated based on the rate of movement of the ink.

### Triglyceride and cholesterol measurement

For whole fly bodies, single fly was anesthetized and transferred to 400 μL of 0.5% PBST, subject to homogenization (Tissuelyser 24, Qiagen, USA) and incubation at 92°C for 10 min. After centrifugation, the supernatant was measured by Triglyceride Quantification Colorimetric/ Fluorometric Kit (BioVision, USA).

For fly hemolymph, extracted hemolymph was directly measured by Triglyceride Assay or Cholesterol Assay Kit (Nanjing Jiancheng Bioengineering Institute, China).

### Hemolymph extraction

40 flies were decapitated and transferred to a punctured 0.5 ml tube. The tube was then placed into 1.5 ml eppendorf tube, subjected to centrifugation for 5 min at 2500 × *g* at 4 °C. Collected hemolymph in 1.5 ml eppendorf tube was used for further analysis.

### LC-MS/MS

LC-MS/MS analysis was performed on HPLC chromatography system named Thermo Fisher Easy-nLC 1000 equipped with a C18 colume (1.8 mm, 0.15×1. 00 mm). The MS/MS data were searched with MaxQuant (version 1.5.2.8). The following parameters were used: (i) enzyme: trypsin; (ii) fixed modification: carbamidomethyl (C); (iii) variable modifications: oxidation (M) and deamidation (NQ); (iv) mass tolerance for precursor ions: 20 ppm; (v) mass tolerance for fragment ions: 4.5 ppm; (vi) peptide charge: 1+, 2+ and 3+; (vii) instrument: ESI-LTQ; (viii) allowing up to two missed cleavage. (ix) minimal peptide length: sixe amino acid. (x) the false discovery rate (FDR) for peptide and protein identifications: 0.01. The intensity for each protein was obtained from three technical repeats. The global analysis was carried by in-house program. The criteria for the interesting proteins were: the ratio of intensity between the experimental and control samples larger than 2 and p-value < 0.05 was considered to be statistically significant.

### Western blotting and Dot blotting

For western blot, total protein was extracted from fly heads or cultured S2 cells. Samples were denatured, separated by SDS-PAGE, and transferred to a polyvinylidene difluoride membrane. After being blocked in TBST containing 5% milk, the membrane was incubated with the specific primary antibody followed by HRP-conjugated goat-anti-rabbit or goat anti-mouse antibody. The specific bands were detected by an ECL western blotting detection system (Bio-rad, USA).

For dot blot, hemolymph was extracted and spotted onto nitrocellulose membrane and let dry for 20 min. After washing, blocking, incubation with the corresponding primary and secondary antibodies, the membranes were also analyzed by an ECL western blotting detection system.

### Quantitative RT-PCR

Total RNA from fly heads was isolated from 5-day-old female flies with TRIzol reagent (Invitrogen, USA). Reverse transcription of RNA to cDNA was achieved using 2 μg of total RNA with random hexamer primers using RT-PCR SuperMix Kit (TransGen Biotech, China). Q-PCR was performed with SYBR premix Ex TaqTM (Takara, China) on CFX96 Touch™ Real-Time PCR Detection System (Bio-Rad, USA). The primers used are presented as follows: DILP2 F: 5′-GCCTTTGTCCTTCATCTCG-3′, DILP2 R: 5′-CCATACTCAGCACCTCGTTG-3′; AKH F: 5′-ATCCCAAGAGCGAAGTCC-3′, AKH R: 5′-CCTGAGATTGCACGAAGC-3′; AKHR F: 5′-ACTGCTACGGAGCCATTT-3′, AKHR R: 5′-TGTCCAGCCAGTACCACA-3′; InR F: 5′-GCAGCAATGAATGCGACGAT-3′, InR R: 5′-CCTGCGTCGCTTGTTGAAAA-3′; GAPDH F: 5′-GAATCCTGGGCTACACCG-3′, GAPDH R: 5′-CTTATCGTTCAGAATGC-3′.

### Statistical analysis

Data presented in this study were verified for normal distribution by D’Agostino–Pearson omnibus test. Student’s t test, one-way ANOVA and two-way ANOVA (for comparisons among three or more groups and comparisons with more than one variant) were used. The post hoc test with Bonferroni correction was performed for multiple comparisons following ANOVA.

## ACKNOWLEDGEMENTS

We thank members of the Neuroscience Pioneer Club for insightful discussions throughout the course of the study. We thank all Wang Lab members and Chen Pan (Zhejiang University) for helpful discussions and technical assistance. We thank Fuchou Tang (Peking University) for help with the single cell analysis. We thank Yeguang Chen (Tsinghua University) and Wei Liu (Zhejiang University) for helpful comments. Ye Wu provide scientific and administrative support in the laboratory. This study was funded by the National Natural Science Foundation of China (No. 31522026 for L.W. and No. 31800883 for R.H.), the Thousand Young Talents Plan (L.W.), the Fundamental Research Funds for the Central Universities (No. 2016QN81010 and No. 2015XZZX004-33 for L.W.), and the 100 Talents Program of Chongqing University (R.H.).

---

**Figure 1-figure supplement 1.**
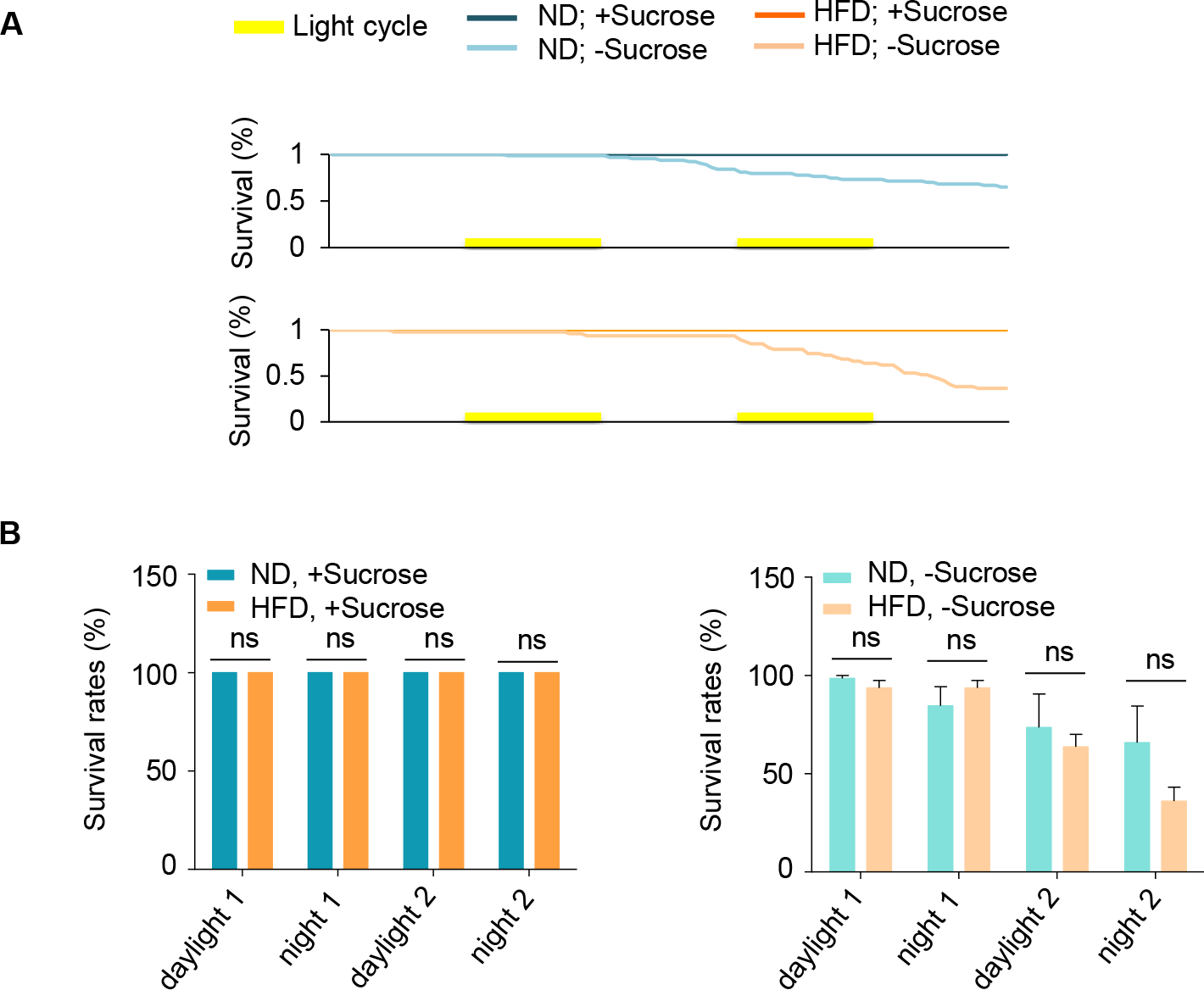
HFD-feeding does not affect flies’ survival. (A) The survival curves of Wild-type *Oregon-R* virgin female flies fed with normal fly food (ND, blue) or high-fat diet (HFD, orange) for five days and then assayed in the presence (dark color) and absence (light color) of 5% sucrose using DAMS-based locomotion assay (n = 46-63). (B) The percentage of live flies at different time points. ns, p > 0.05; *p < 0.05; ***p < 0.001; ****p < 0.0001. Student’s t-test and one-way ANOVA followed by post hoc test weith Bonferroni correction were used for pair-wise and multiple comparisons, respectively.

**Figure 4-figure supplement 1.**
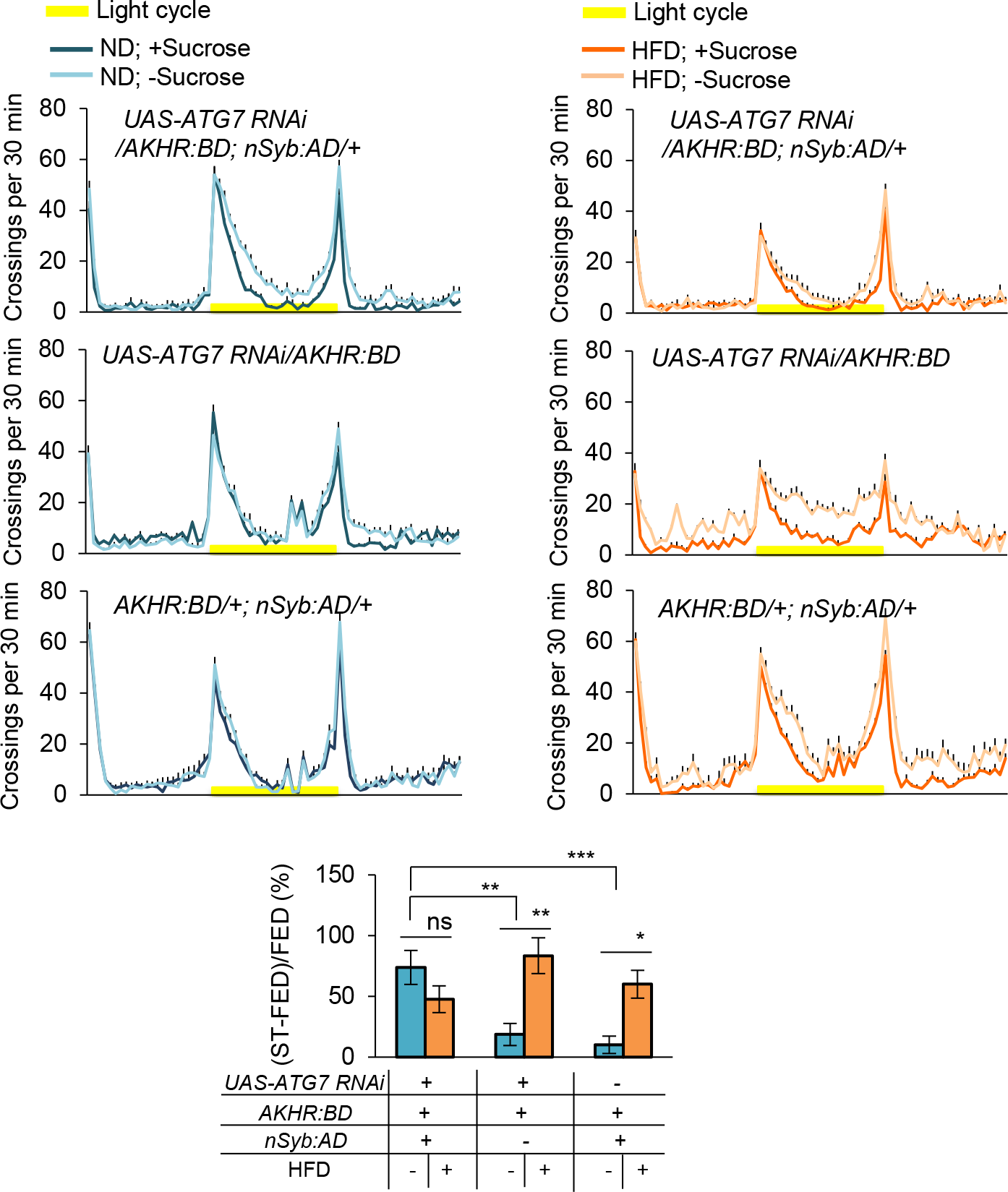
Knocking down ATG7 in AKHR^+^ neurons promotes starvation-induced hyperactivity. Midline crossing activity of indicated genotypes assayed in the presence or absence of 5% sucrose in Day 1 (upper) (n = 30-60). Starvation-induced hyperactivity after 5-day HFD feeding vs. ND feeding experience is also shown (lower). ns, p > 0.05; *p < 0.05; ***p < 0.001; ****p < 0.0001. Student’s t-test and one-way ANOVA followed by post hoc test weith Bonferroni correction were used for pair-wise and multiple comparisons, respectively.

**Figure 5-figure supplement 1.**
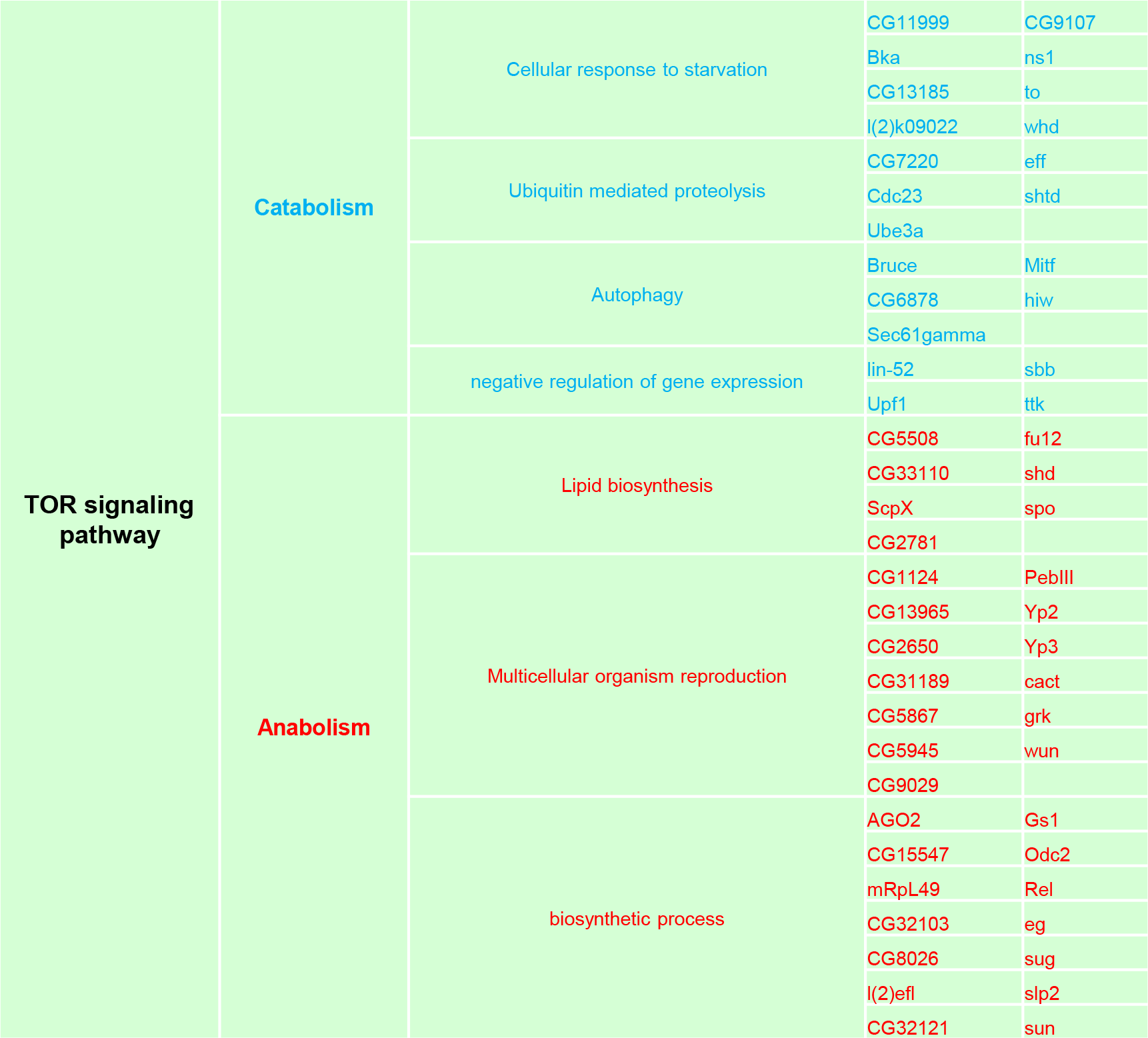
TOR signaling pathway-related genes that are differentially expressed in the head tissues of flies fed with ND vs. HFD. Wild-type virgin female flies were fed with ND or HFD. Their head tissues were harvested and subjected to RNAseq analysis. GO categories and KEGG pathway analysis were used to identify TOR signaling pathway-related genes, whose expression levels were further analyzed in Figure 5-figure supplement 2. Red and blue colors indicate anabolism and catabolism genes, respectively.

**Figure 5-figure supplement 2.**
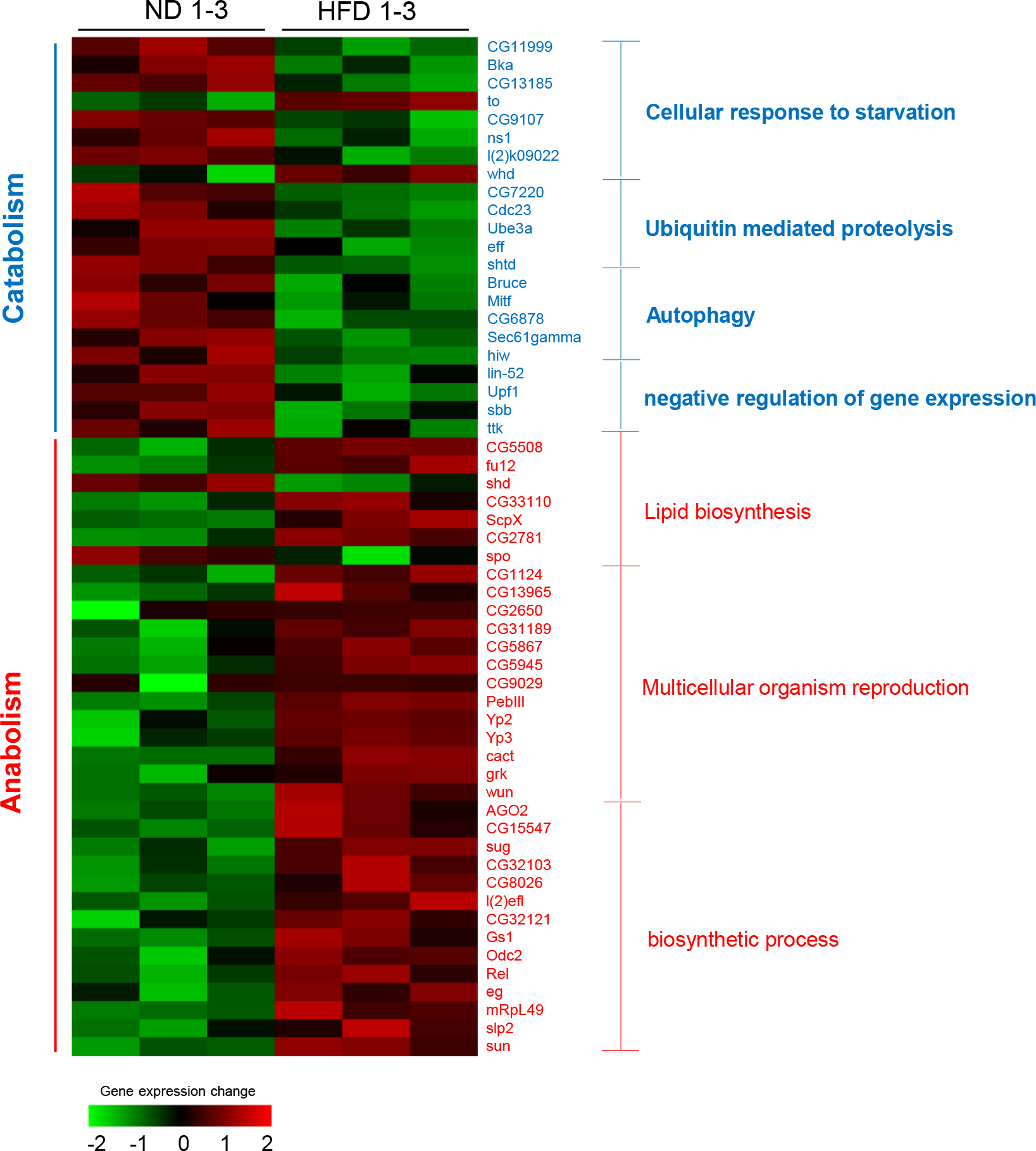
Differentially expressed genes in TOR signaling pathways in the head tissues of flies fed with ND vs. HFD. Heat map of differentially expressed genes enriched in TOR signaling pathways. Wild-type virgin female flies were fed with ND or HFD. Their head tissues were harvested and subjected to RNAseq analysis. Green and red columns represent down-regulated genes and up-regulated genes, respectively. Red and blue font colors indicate anabolism and catabolism genes, respectively. Note that most anabolism related genes were up-regulated while most catabolism related genes were down-regulated upon HFD feeding. See Figue 5-figure supplement 1 for the identification/classification of these genes.

**Figure 6-figure supplement 1.**
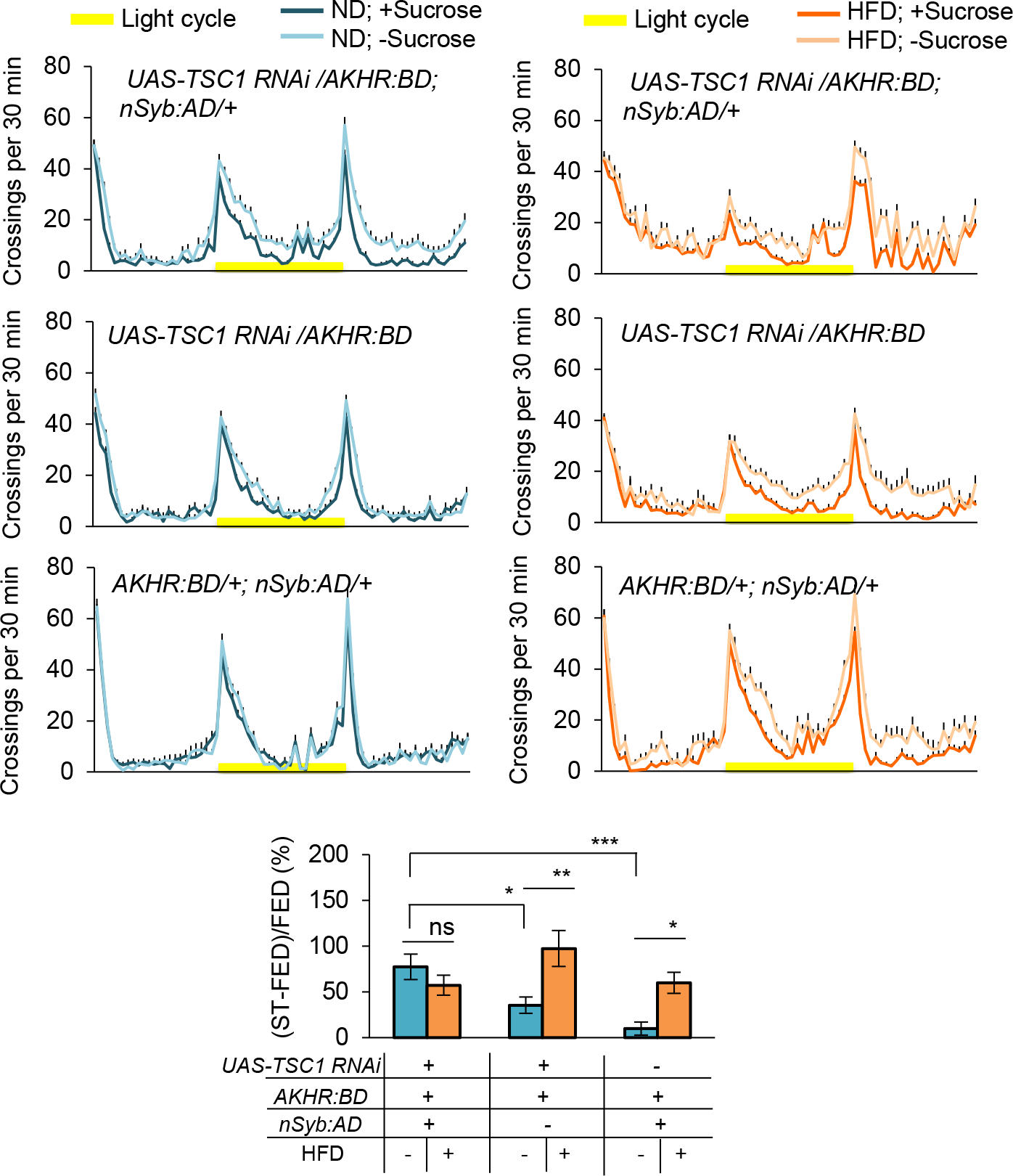
Activation of TOR signaling in AKHR^+^ neurons promotes starvation-induced hyperactivity. Midline crossing activity of indicated genotypes assayed in the presence or absence of 5% sucrose in Day 1 (upper) (n = 36-46). Starvation-induced hyperactivity after 5-day HFD feeding vs. ND feeding experience is also shown (lower). ns, p > 0.05; *p < 0.05; ***p < 0.001; ****p < 0.0001. Student’s t-test and one-way ANOVA followed by post hoc test weith Bonferroni correction were used for pair-wise and multiple comparisons, respectively.

**Figure 6-figure supplement 2.**
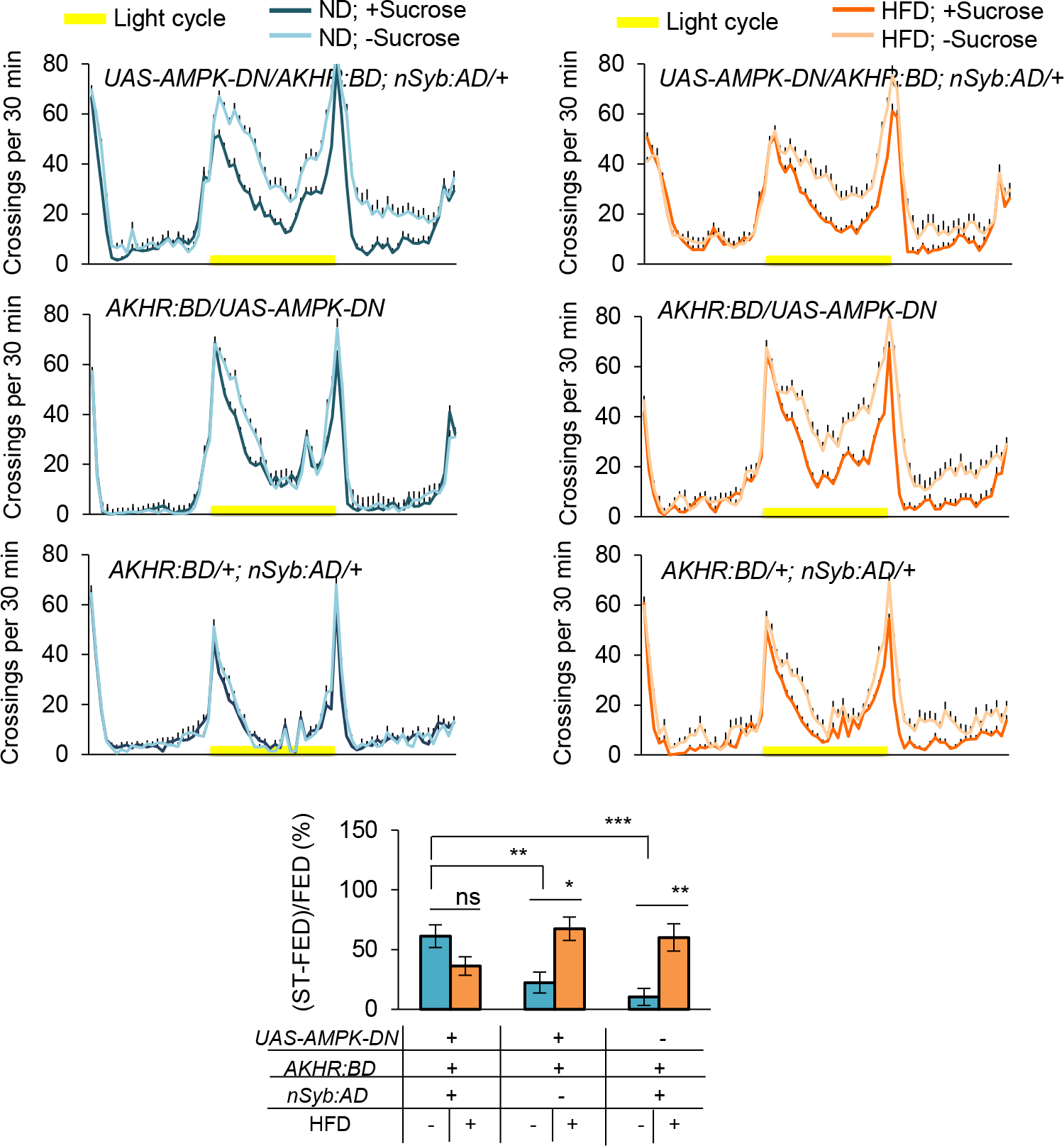
Inhibition of AMPK signaling in AKHR^+^ neurons promotes starvation-induced hyperactivity. Midline crossing activity of indicated genotypes assayed in the presence or absence of 5% sucrose in Day 1 (upper) (n = 40-59). Starvation-induced hyperactivity after 5-day HFD feeding vs. ND feeding experience is also shown (lower). ns, p > 0.05; *p < 0.05; ***p < 0.001; ****p < 0.0001. Student’s t-test and one-way ANOVA followed by post hoc test weith Bonferroni correction were used for pair-wise and multiple comparisons, respectively.

**Figure 8-figure supplement 1.**
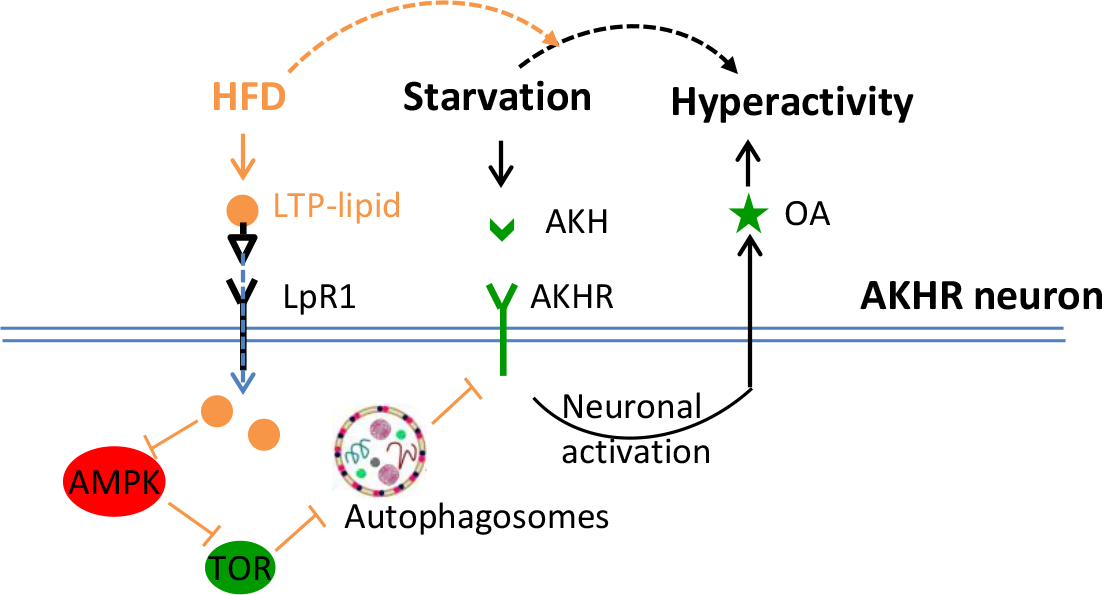
A working model. We propose the following working model based on our results. AKHR protein levels in AKHR^+^ neurons are well maintained via protein expression and autophagic degradation. Under ND feeding, starvation induces the expression and release of AKH, which in turn activates AKHR^+^ neurons and hence the release of OA. OA signaling promotes starvation-induced hyperactivity in these starved flies. Upon HFD feeding, however, the regulation of starvation-induced hyperactivity is disrupted and greatly enhanced. Mechanistically, LTP delivers excess dietary lipids to AKHR^+^ neurons via its cognate receptor LpR1. Subsequently, neuronal autophagy is suppressed in AKHR^+^ neurons via AMPK-TOR signaling, which leads to AKHR accumulation. As a result, AKHR^+^ neurons become hyper sensitive to AKH and therefore starvation-induced hyperactivity is enhanced.

